# Cysteine availability tunes ubiquitin signaling via inverse stability of LRRC58 E3 ligase and its substrate CDO1

**DOI:** 10.1101/2025.11.14.688510

**Authors:** Gisele A. Andree, Luca Stier, Kerstin Schmiederer, Alina Thielen, Luis Schmid, Samuel A. Maiwald, Jiale Du, Susanne von Gronau, Claudia Strasser, Judith Müller, Lukas T. Henneberg, Camille Guyot, Gary Kleiger, Matthias Mann, Peter J. Murray, Brenda A. Schulman

## Abstract

Cellular responses to amino acid fluctuations often hinge on ubiquitin-mediated control of metabolic enzymes, yet the underlying E3 ligase pathways remain incompletely defined. Using quantitative proteomics and active cullin-RING ligase (CRL) profiling, we identify LRRC58 as a cysteine-responsive substrate receptor whose stability increases sharply under cysteine starvation. Proteomics reveals an inverse relationship between LRRC58 and the metabolic enzyme cysteine dioxygenase 1 (CDO1), suggesting a cysteine-linked regulatory axis. Biochemical reconstitution and cryo-EM structures show that LRRC58 forms an active CUL2-or CUL5-based CRL that selectively positions CDO1 for ubiquitylation at Lys8. By contrast, targeted protein degradation via the VHL-based CRL is achieved by recruitment of a distinct surface of CDO1, and more promiscuous ubiquitylation independent of Lys8. Together, our proteomics-guided discovery pipeline, cellular stability studies, and structural analyses uncover a metabolically-tuned LRRC58-CDO1 pathway that links cysteine availability to selective proteasomal turnover, reveals principles of metabolite-regulated CRL activity, and showcases mechanisms distinguishing endogenous and targeted protein degradation.

## Introduction

Regulation of metabolic enzyme levels is critical for maintaining cellular homeostasis in different conditions, such as nutrient limitation or excess, and in response to availability of essential cofactors and substrates. It has been known for over half a century, that enzyme concentrations are not only determined by new protein synthesis, but also by specific degradation pathways responding to changing metabolic cues^1^. In eukaryotes, the ubiquitin-proteasome-system is often employed to target specific proteins for degradation^2–4^. Central to the UPS are E3 ubiquitin ligases, which provide specificity through selectively binding substrates and marking them with ubiquitin signals for degradation.

In human cells, E3 ligases and their cognate substrates have been identified as coordinately responding to metabolic signals including carbohydrates, lipids, ions, metabolic cofactors, and redox stresses^5–14^. Additional modes of regulation have been observed for dipeptides controlling amino acid uptake in yeast and lipid homeostasis in human cells^15,16^. Furthermore, some metabolites control the levels of the enzymes catalyzing their biosynthesis and catabolism. An archetypal example of E3 ligase-dependent control of metabolic pathways is the multiprotein budding yeast “GID” E3 ligase complex, discovered in screens for mutants that were “Glucose Induced Degradation Deficient” for a Fructose-1,6-bisphosphatase reporter system^17–22^. The substrate binding subunit of the GID E3 complex is induced by glucose, driving ubiquitin-mediated degradation of its gluconeogenic enzyme substrates under conditions when their activities become superfluous^23,24^. Similarly, when sterol levels are high, they bind SQLE and HMGCR, key enzymes in the cholesterol biosynthetic pathway, to induce their ubiquitylation and degradation^25–34^.

Amino acids also affect the stabilities of enzymes that catalyze reactions pertaining to their metabolism^1^. Importantly, cysteine homeostasis has long been known to rely on regulation by the ubiquitin-proteasome pathway. For example, cysteine availability controls the stability of cysteine dioxygenase type 1 (CDO1), which catalyzes oxygenation of cysteine for biosynthesis of hypotaurine and taurine. A series of discoveries in rodents revealed that: (1) dietary cysteine controls CDO1 activity; (2) the cellular abundances of cysteine and CDO1 protein are correlated; (3) limiting cysteine availability triggers degradation of CDO1 by the proteasome; and (4) CDO1 is stable when cysteine is replete^35–37^. Until recently however, how CDO1 is directed for degradation has been a mystery, at least in part due to lack of knowledge of its specific E3 ligase(s).

To discover E3 ligases activated and deactivated in response to shifts in environmental conditions, including changes in metabolic state, we developed active cullin-RING ligase (CRL) profiling technology^38^. CRLs are a collection of hundreds of modular E3 complexes, wherein a catalytic cullin-RING module binds a substrate-binding receptor^39,40^. Most CRLs contain either the RING protein RBX1 partnered with CUL1, CUL2, CUL3, or CUL4, or the similar RBX2-CUL5 complex. Each cullin-RING module binds interchangeably to numerous substrate-binding receptors. It is the substrate receptor that specifies the target for ubiquitylation. CRLs are activated by post-translational modification of the cullin subunit with the ubiquitin-like protein NEDD8^41,42^. A neddylated cullin and its partner RING protein recruit and activate a ubiquitin carrying enzyme (an E2 or ARIH-family RBR E3 ligase) that covalently links ubiquitin to the CRL’s receptor-bound substrate^43–47^.

Neddylation status of a particular CRL is tightly regulated in accordance with cellular demand for that E3’s activity^48–51^. As such, active CRL profiling, which applies quantitative proteomics to affinity-enriched neddylated CRLs, can identify specific substrate receptors whose association with neddylated cullins are modulated in response to changes in metabolic conditions^38^.

To discover E3s regulated based on cysteine abundance, we applied active CRL profiling. Assaying multiple cell lines in parallel revealed that cysteine starvation activates a CRL with the substrate receptor, LRRC58. Cellular stability studies revealed that LRRC58 targets CDO1 for degradation in cysteine limiting conditions, and that LRRC58 is destabilized and CDO1 stabilized when its cysteine substrate is abundant. This pathway was independently reported by others using different methodologies while our manuscript was in preparation^52–54^. Our biochemical reconstitution and cryo-EM, in comparison with the targeted protein degradation of CDO1 by a different CRL^55^, revealed unique CDO1 ubiquitylation by LRRC58.

## Results

### LRRC58-CRL is activated upon cysteine starvation

We applied active CRL profiling to lysates from HeLa cells cultured for 24 hours in either normal or cysteine-free media, or cultured in normal media for 30, 60, or 120 minutes following 24 hours of cysteine starvation. Immunoprecipitations (IP) performed with our antigen-binding fragment (Fab) that specifically recognizes neddylated CUL1, CUL2, CUL3 and CUL4 (but not other cullins) were analyzed by high-resolution mass-spectrometry-based proteomics using data-independent acquisition (DIA-MS) (Fig. 1a). When comparing the samples from 24 hour cysteine starvation to the control, only one CRL substrate receptor was enriched: the BC-box protein LRRC58 (Fig. 1b). LRRC58 remained enriched after exchange into normal media. Based on these initial experiments, we also examined the active CRLs from HEK293T cells that vary upon a shift in cysteine amounts. Here, the comparison was between cells grown for 24 hours in cysteine-free or in normal media. LRRC58 was the only CRL substrate receptor identified in cysteine starvation samples that was not detected in samples from growth in normal media. Furthermore, LRRC58 displayed the highest normalized intensity of all proteins that were undetectable when cells were grown on normal media (Fig. 1c).

**Figure 1.**
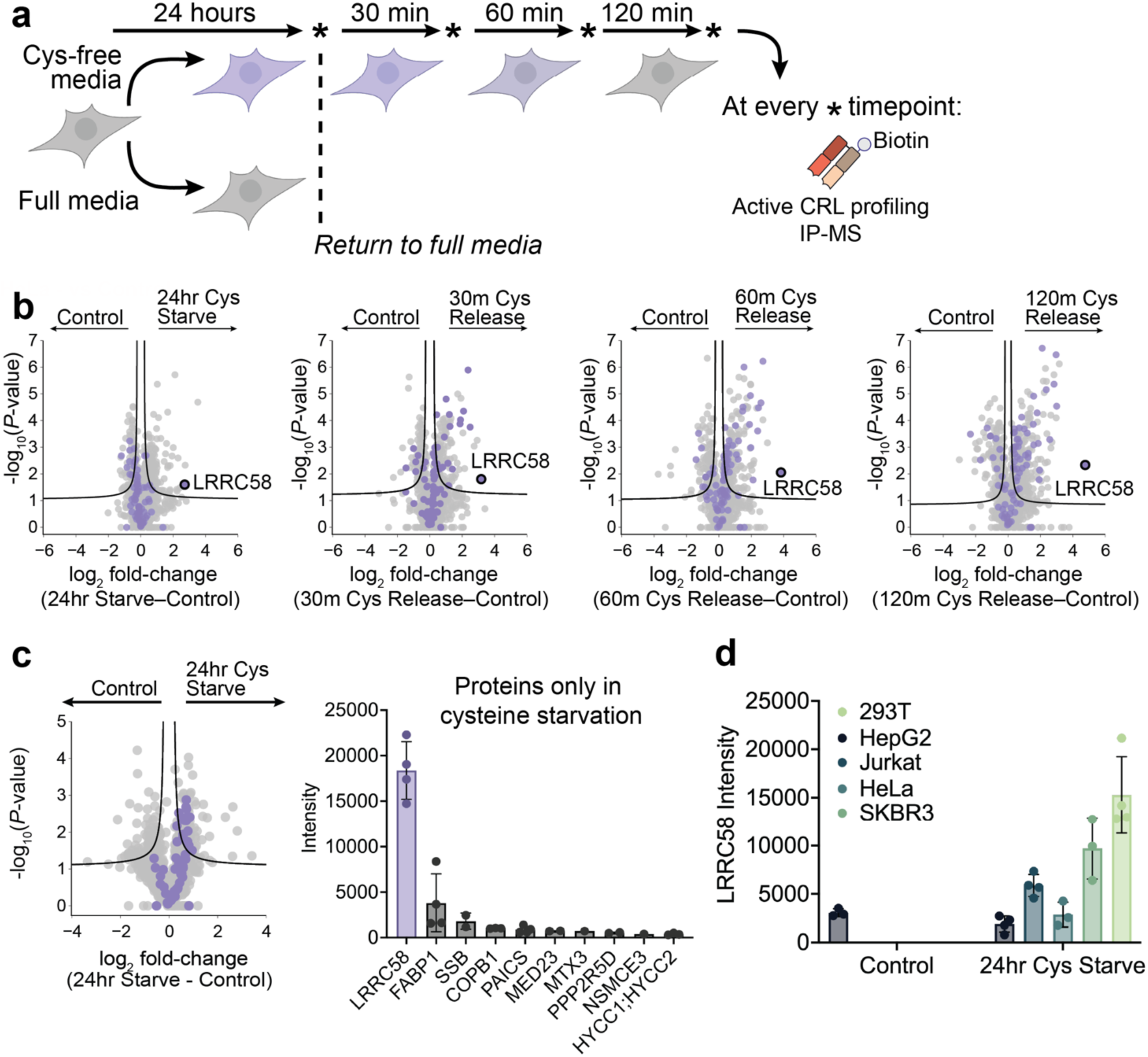
Protein levels increase for the CRL substrate receptor LRRC58 during cysteine starvation. (a) Schematic showing a protocol for cysteine starvation of tissue culture cells and time-dependent return to normal media. CRL substrate receptor levels were assessed through active CRL profiling. (b) Volcano plots highlighting changes in protein levels comparing HeLa cells grown in complete media (control) versus cells grown in cysteine-free media for 24 hours prior to switching to complete media at the indicated time points (*n* = 3 independent replicates). Known CRL substrate receptors are colored in purple, all other identified proteins are in grey. Curves for 5% FDR thresholds are shown (two-sided t test, permutation-based FDR calculation, 0.05 FDR, 250 randomizations, s0=0.1). (c) Volcano plot (left) same as in (b) but with HEK293T cells grown in complete or in cysteine-free media for 24 hours (*n* = 4 independent replicates). Bar graph (right) showing the average intensities of the proteins identified in the IP-MS samples during cysteine starvation that were not detected when cells were grown in complete media. (d) Average LRRC58 intensities from total proteomes of the indicated cell lines grown in either complete (control) or in cysteine-free media for 24 hours (*n* = 4 independent replicates). All error bars report the standard deviation of the data points.

Two broad mechanisms have been found to regulate substrate receptor incorporation into a neddylated CRL. CRLs are often regulated through cycles of deneddylation and neddylation, where deneddylated cullin-RING complexes are subject to additional cycles of disassembly and assembly with their repertoire of substrate receptors. In such cases, substrate binding to the receptor inhibits deneddylation and disassembly, shifting the equilibrium of that substrate receptor towards neddylated complexes^48–50^. The abundance of the substrate receptor, often determined by its own ubiquitin mediated proteolysis, impacts the extent of its incorporation into an active E3^38,56–58^. As a first step towards uncovering how cysteine regulates LRRC58, DIA-MS experiments were performed to quantify the proteomes of five human cell lines (HEK293T – embryonic kidney, HepG2 – liver tumor, Jurkat – T-ALL, HeLa – cervical cancer; SKBR3 – breast cancer) grown for 24 hours in either normal or cysteine-free media. Overall, the amounts of LRRC58 change to a striking extent in response to the level of cysteine in the media (Fig. 1d).

### Cellular LRRC58 and CDO1 protein abundance are inversely correlated in a cysteine-dependent manner

To determine if the increased amount of LRRC58 in cysteine starvation conditions is regulated by transcriptional responses, RNA sequencing (RNA-Seq) was performed for HEK293T cells grown for 24 hours in either normal media or cysteine-free media (Fig. 2a). LRRC58 transcript abundances did not significantly differ between the samples, hinting that the response to cysteine starvation may be due to post-transcriptional events. We next examined potential involvement of ubiquitin-mediated degradation by treating cells with either a proteasome or a neddylation inhibitor (MG132 and MLN4924, respectively). In the absence of such inhibitors, LRRC58 was detected by mass spectrometry-based proteomics analysis only in the cysteine starvation condition (notably, we and others^52,54^ have been unable to obtain an antibody recognizing LRRC58) (Fig. 2b). Meanwhile, higher amounts of LRRC58 were detected, even from cells cultured in normal media, upon treatment with MG132 and MLN4924 (Fig. 2b). Furthermore, the data raise the possibility that LRRC58 may be continuously degraded under normal growth conditions, but is stabilized when cysteine is limiting. Indeed, autodegradation is a common mechanism to negatively regulate substrate receptors when their functions are not needed^38,56,57^.

**Figure 2.**
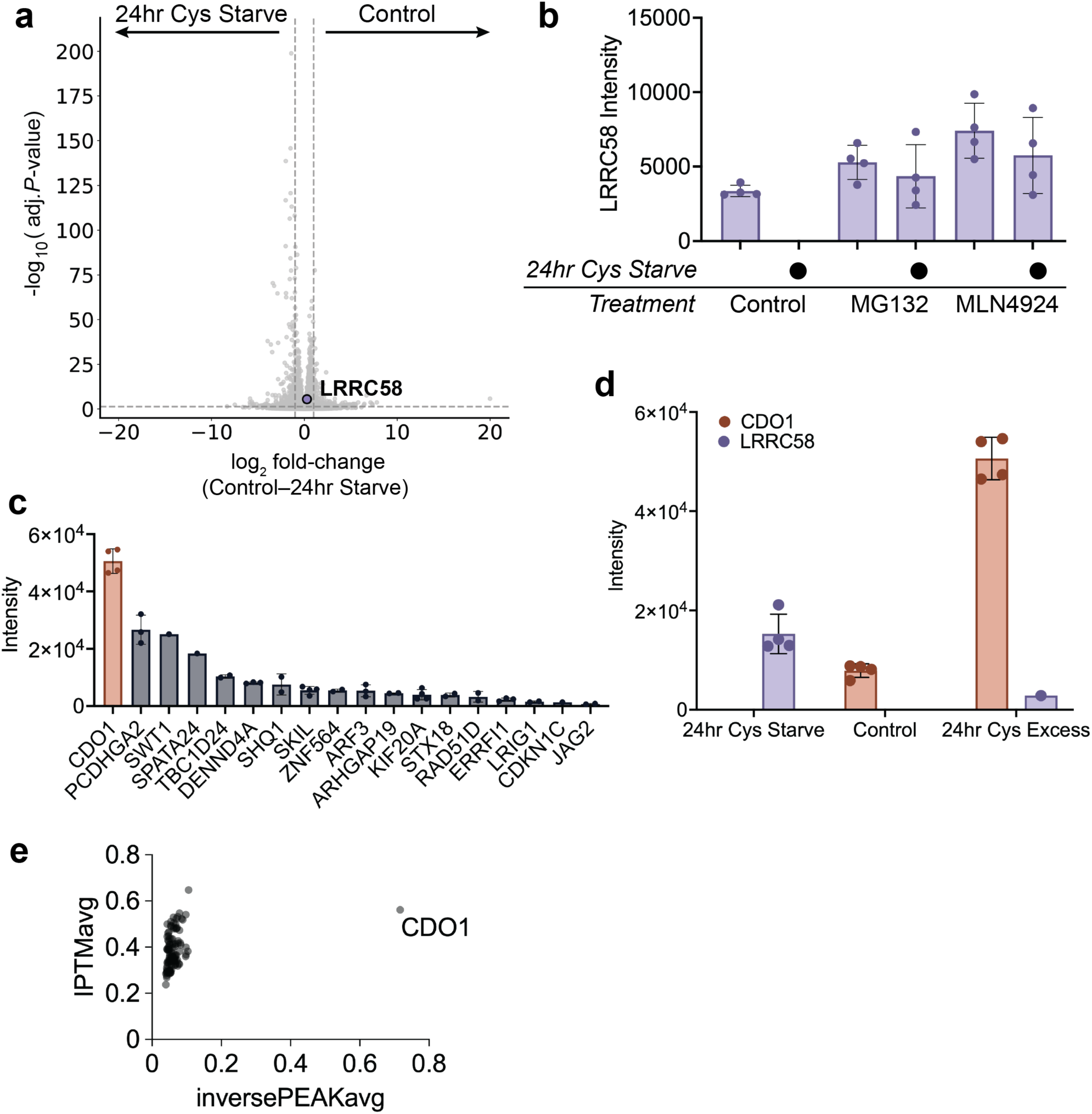
Identification of CDO1 as a putative substrate of LRRC58. (a) Volcano plot from RNA-seq analysis of HEK293T cells grown for 24 hours in either complete or cysteine-free media (*n* = 3 independent replicates). LRRC58 transcript levels (purple) did not change significantly. Vertical and horizontal dashed lines indicate thresholds at a 2-fold change in RNA levels and p-value = 0.05 (adjusted p-values (Benjamini-Hochberg procedure) ≤ 0.01), respectively. (b) Average LRRC58 intensities identified in the total proteomes (*n* = 4 independent replicates) of HEK293T cells treated with proteasome (MG132) or neddylation inhibitors (MLN4924). Cells were grown in either complete or cysteine-free media for 24 hours. (c) Average intensities of proteins identified from the total proteome of HEK293T cells (*n* = 4 independent replicates) which showed detectable levels when grown in the presence of 10-fold excess cysteine but not when grown in cysteine-free media. (d) Average intensities of LRRC58 and CDO1 identified in the total proteomes of HEK293T cells comparing growth after 24 hours in cysteine-free media, complete media (control), or with 10-fold excess cysteine (*n* = 4). CDO1 protein was undetectable in cells grown in cysteine-free media whereas LRRC58 was undetectable in the control. (e) The average IPTM and average inverse PEAK scores calculated from HT-Colabfold analysis of the interactions between LRRC58-EloB/C and all proteins absent in the total proteomes of cysteine starvation samples, but detected upon growth with excess cysteine for the cell lines; HEK293T, HepG2, Jurkat, HeLa, and SKBR3. All error bars report the standard deviation of the data points.

We reasoned that proteins with levels that are lower when LRRC58 is expressed (i.e. in the cysteine starvation conditions), but that accumulate in the presence of excess cysteine when LRRC58 abundance is low could represent putative substrates of this E3. We thus performed DIA-MS to quantify proteins from HEK293T cells grown in the absence of cysteine with those cultured with a 10-fold excess cysteine concentration. The most abundant protein that increased in the presence of cysteine was CDO1 (Fig. 2c). This observation is consistent with the known regulation of CDO1 that depends on cysteine availability^35–37^. Importantly, CDO1 and LRRC58 protein levels were inversely correlated. Peptides corresponding to CDO1 were not detected from the cysteine starved sample, for which LRRC58 abundance was high (Fig. 2d).

An additional four cell lines (HepG2, Jurkat, HeLa, SKBR3, also studied in Fig. 1d) were treated and analyzed similarly. Proteins absent in the cysteine starvation conditions, but detected upon growth with excess cysteine were considered potential LRRC58 substrates. All such proteins arising from any of the cell lines were modeled for potential interaction with LRRC58 (and its obligate CRL partners Elongin B and Elongin C, hereafter EloB/C) using HT-Colabfold^59^. Notably, only CDO1 scored as a potential interactor with the LRRC58 complex (Fig. 2e).

### CDO1 stability reporter responds to cysteine starvation in an LRRC58-and CUL2-dependent manner

In order to probe the regulation of CDO1 degradation, we used a stability reporter system^60^. CDO1 was expressed as an N-terminal fusion to a fluorescent mCherry-tag in a dual-fluorophore reporter construct (GFP–P2A–mCherry-CDO1) (Fig. 3a, Extended Data Fig. 1a). Reporter-expressing cells are GFP-positive while the fraction of cells lacking mCherry fluorescence reflects CDO1 degradation. To validate this system, we first confirmed that the CDO1 reporter reflects the cysteine-dependent regulation determined for endogenous proteins by DIA-MS. Indeed, the percentage of mCherry-negative cells agrees with the known regulation: low for growth in cysteine, high for growth in the absence of cysteine (Fig. 3a). Second, we confirmed that destabilization of the CDO1 reporter depends on a cullin-RING E3 ligase. The percentage of mCherry-negative cells is low even for cysteine starvation when cells were treated with neddylation inhibitor (MLN4924). The reporter also showed stabilization upon proteasome inhibition (MG132) (Fig. 3b). Finally, siRNA-mediated knockdown of LRRC58 also stabilized the CDO1 reporter (Fig. 3c, Extended Data Fig. 1b). Taken together, these data support our experimental validity of using a stability reporter as a surrogate readout for the LRRC58-CRL induced degradation of CDO1.

**Figure 3.**
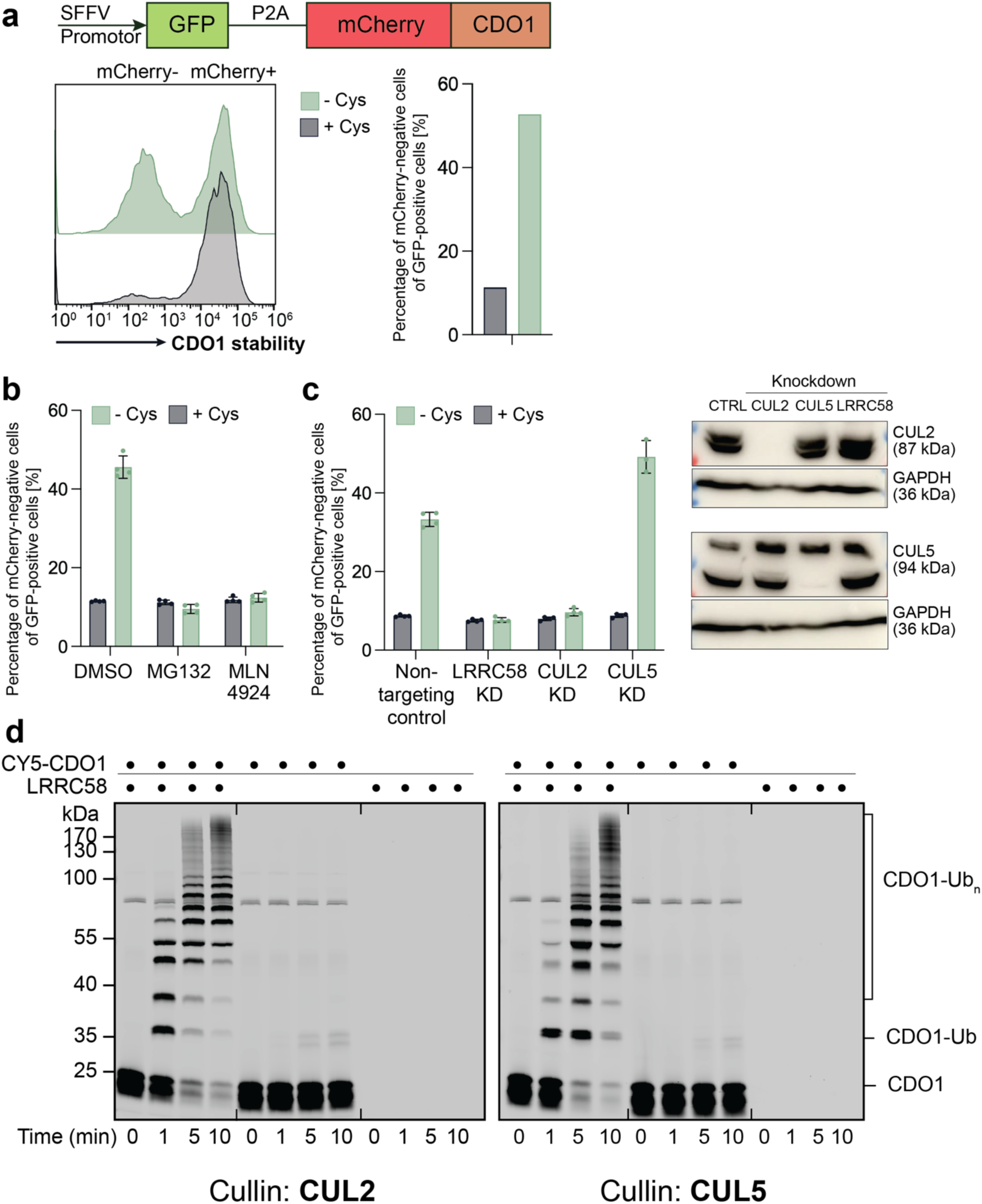
Cysteine depletion reduces cellular CDO1 levels in a CUL2-dependent manner. (a) The CDO1 reporter is a dual-fluorophore reporter construct with GFP and CDO1 N-terminally tagged with mCherry separated by a P2A sequence. The CDO1 reporter is monitored by flow cytometry in the presence and absence of extracellular cysteine. A low percentage of mCherry-negative cells (of all GFP positive cells) reflects CDO1 stabilization while a higher percentage of mCherry-negative cells (of all GFP positive cells) indicates CDO1 destabilization, as displayed in the bar graph on the right. (b) Reporter assay showing decreased CDO1 levels in response to cysteine starvation. CDO1 downregulation was sensitive to CRL and 26S proteasome activities (MLN4924 and MG132 treatments, respectively). Bars represent the average values from n=4 independent replicates. (c) Decrease in CDO1 reporter levels upon cysteine starvation is dependent on the presence of LRRC58 and CUL2 but not CUL5. HEK293T cells were treated with siRNAs targeting LRRC58, CUL2, or CUL5 expression or non-targeting control (n=4 independent replicates). Representative Western blot (right) showing the efficiency of si-RNA mediated CUL2 and CUL5 knockdown (KD). LRRC58 KD efficiency was determined through analysis of the total proteome (Extended Data Fig. 1b). All error bars report the standard deviation of the data points. (d) In vitro reconstituted ubiquitylation of Cy5-labeled CDO1 in the presence of neddylated CUL2-RBX1 or CUL5-RBX2 and in the absence or presence of LRRC58-EloB/C. Fluorescence scans are representative of n=3 technical replicates.

With the validated reporter in hand, we next sought to discover a cullin component mediating CDO1 degradation. LRRC58 has been annotated as a BC-box protein binding to EloB/C^61^. EloB/C can recruit BC-box substrate receptors to CUL2 or CUL5. Since our current active CRL profiling excludes CUL5^38^, our data suggested LRRC58 could associate with CUL2-RBX1 to form an active E3 (Fig. 1). Published analyses of the amino acid sequences of BC-boxes predicted LRRC58 association with CUL2^61^. However, previous high-throughput interactome studies showed LRRC58 binding to CUL5^62–64^.

While the siRNA-mediated knockdown of CUL2 or CUL5 protein levels were robust (Fig. 3c), significant CDO1 reporter stabilization was observed only upon knockdown of CUL2. Similar data were posted on bioRxiv during our manuscript preparation^52^. That study additionally found that the co-silencing of *CUL5* with *CUL2* led to maximal stabilization of a similar CDO1 reporter.

### CDO1 binds to LRRC58 and is ubiquitylated by activated CRL complexes in vitro

To further define the basis for LRRC58 regulation of CDO1, we reconstituted interactions with purified components (Extended Data Fig. 2). After mixing purified CDO1 with an LRRC58-EloB/C complex, we observed co-migration by size-exclusion chromatography indicative of a stoichiometric assembly. Similarly, addition of CUL2-RBX1 led to formation of a CRL-substrate complex. A parallel complex was also formed with CUL5-RBX2. Both CRLs ubiquitylated CDO1 in an LRRC58-dependent manner in biochemical assays (Fig. 3d). Hereafter, we refer to the CRL complexes with the different cullins as LRRC58-CUL2 and LRRC58-CUL5.

### Distinct CDO1 targeting by endogenous versus molecular glue-induced degradation machineries

CDO1 was recently identified as the target of molecular glue degraders hijacking the CUL2 substrate receptor VHL^55^. We investigated mechanistic differences for CDO1 ubiquitylation through its natural substrate receptor LRRC58, and VHL together with the small molecule “compound-8” (abbreviated “Cmpd8”). Consistent with the literature, the addition of compound-8 to cells destabilized the CDO1 reporter (Fig. 4a). We also compared the two mechanisms of CDO1 substrate recruitment using our reconstituted in vitro ubiquitylation assay (Extended Data Fig. 3). These biochemical experiments varied only by substrate receptor, and employed CUL2-RBX1 that is utilized by both E3s. In vitro ubiquitylation of CDO1 by the VHL-CUL2 neddylated CRL depended on compound-8, and vice-versa.

**Figure 4.**
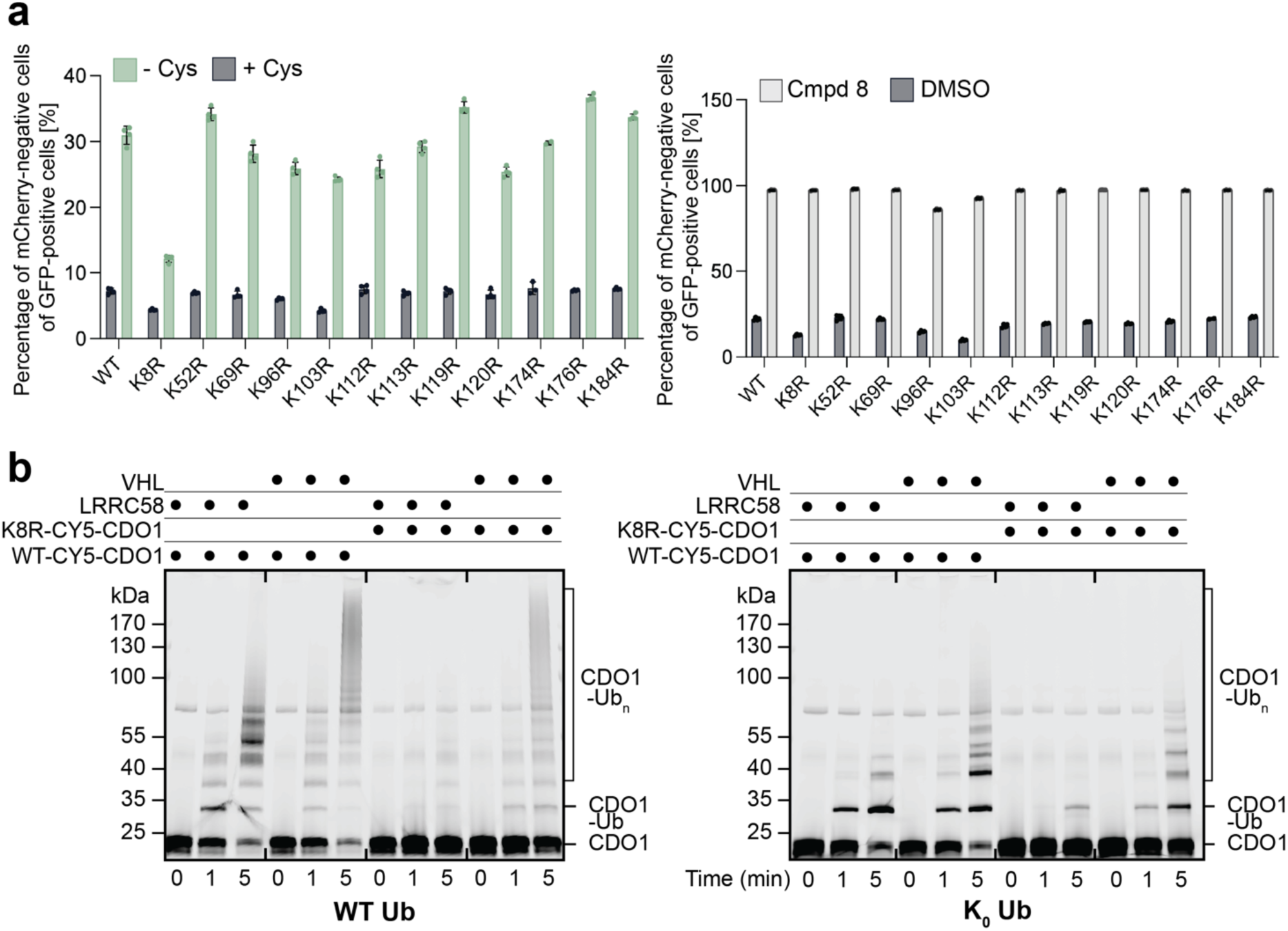
Selective CDO1 Lys targeting during native ubiquitylation is reprogrammed into promiscuous targeting by a molecular glue. (a) Left panel displays the reporter assay showing single Lys to Arg CDO1 reporter variants expressed in HEK293T cells grown in complete or cysteine-free media. K8R CDO1 levels were minimally affected upon cysteine starvation in contrast to all other CDO1 mutants, which showed wild-type like responses. Right panel is the same as the left except comparing cells gown in complete media and treated with Compound 8 (Cmpd 8) or DMSO. All CDO1 reporter variants are efficiently degraded. Bars represent the average values from n=4 independent replicates. Error bars report the standard deviation of the data points. (b) Left panel shows in vitro reconstituted assays comparing Cy5-labeled WT and K8R CDO1 ubiquitylation in the presence of neddylated LRRC58-CUL2 or neddylated VHL-CUL2 with Cmpd8. The efficiency of K8R CDO1 ubiquitylation is significantly lower with LRRC58-CUL2 compared to WT CDO1 but minimally affected in the presence of VHL-CUL2 and Cmpd8. Assay performed with wild-type ubiquitin (WT Ub). Right panel is the same as left but performed using a lysine-less ubiquitin (K_0_-Ub) that cannot form chains.

Both cellular and in vitro ubiquitylation assays (Fig. 4, Extended Data Fig. 3) surprisingly showed that CDO1 targeting was less efficient through LRRC58 than by targeted protein degradation. Based on prior studies examining substrate targeting^44–47,65,66^, we hypothesized that the native degradation mechanism could be constrained if one or more specific lysines are selectively targeted. Thus, we tested effects of arginine replacements (which cannot accept ubiquitins) for individual lysines in the CDO1 stability reporter (Fig. 4a). Strikingly, a single K8R substitution was impaired for cysteine-dependent destabilization of CDO1. However, all CDO1 variants were readily destabilized by compound-8.

These data raised the possibility that Lys8 is preferentially targeted by the LRRC58 E3. Indeed, in ubiquitylation assays using the LRRC58 CRL, there was little modification of the K8R mutant compared to WT CDO1. This defect was observed in assays using both WT ubiquitin and a lysine-less version (K0-ubiquitin) that cannot form chains. The data suggest Lys8 modification serves as a prerequisite for chain formation in the LRRC58 pathway. Notably, the compound 8-VHL system retained activity towards the K8R mutant CDO1 (Fig. 4b). Overall, our experimental findings reveal that targeted protein degradation approaches can lead to superior substrate turnover by circumventing limitations imposed by native modes of substrate engagement, and that CDO1 is distinctly presented to the CRL catalytic machinery by its endogenous substrate receptor LRRC58.

### Cryo-EM reconstructions visualizing CDO1 ubiquitylation by LRRC58 CRL

To understand the strikingly specific ubiquitylation on CDO1 Lys8 by LRRC58, we sought structural data, by cryo-EM. We generated samples for both CUL2 and CUL5-based CRLs in parallel, considering that (1) structure determination is an empirical process where unpredictable protein factors can affect data quality, and (2) LRRC58 can engage both these cullins^62–64^ (Fig. 3c,d, Extended Data Fig. 2). We generated stable mimics of ubiquitylation intermediates applying our established chemical biology method^44,46,47,67,68^. Our approach simultaneously links the active site of a ubiquitylating enzyme, a modified C-terminus of ubiquitin, and a cysteine-substitution for the targeted Lys (here CDO1 Lys8). We used ARIH1 as the ubiquitylating enzyme with neddylated CUL2-RBX1, and ARIH2 with neddylated CUL5-RBX2. Previous studies have shown that ARIH-family enzymes broadly and efficiently target CRL substrates with a wide range of structural features^43,44,69^. Cryo-EM reconstructions visualized LRRC58 recruitment of CDO1 for ubiquitylation, at 3.73 Å and 2.95 Å overall resolution for neddylated CUL2-RBX1-ARIH1 and CUL5-RBX2-ARIH2 complexes, respectively (Table 1, Fig. 5a, Extended Data Figs. 4-6).

**Table 1:** Data collection, refinement and validation statistics

|  | Structure representing CDO1 ubiquitylation by LRRRC58 with neddylated CUL5-RBX2-ARIH2 | Screening dataset map representing CDO1 ubiquitylation by LRRRC58 with neddylated CUL2-RBX1-ARIH1 | Maps representing CDO1 ubiquitylation by LRRRC58 with neddylated CUL2-RBX1-ARIH1 |
| --- | --- | --- | --- |
| **Data collection and processing** |  |  |  |
| Magnification | 105,000 | 22,000 | 105,000 |
| Voltage (kV) | 300 | 200 | 300 |
| Electron exposure (e-/Å²) | 50.2 | 60.0 | 51.0 |
| Defocus range (µm) | −0.6 to −2.2 | −1 to −2.6 | −0.6 to −2.2 |
| Pixel size (Å) | 0.8512 | 1.841 | 0.8512 |
| Symmetry imposed | C1 | C1 | C1 |
| Initial particle images (no.) | 17,986,650 | 11,643,225 | 13,861,287 |
| Final particle images (no.) | 847,022 | 206,311 | 1,462,967 |
| Map resolution (Å) | 2.95 | 4.88 | 3.70 |
| FSC threshold | 0.143 | 0.143 | 0.143 |
| Map resolution range (Å) | 2.31 – 8.37 | 4.40 –11.96 | 2.90 – 8.43 |
| **Refinement** |  |  |  |
| Initial model used (PDB code) | AlphaFold3, 4N9F, XXXX, 7B5M, |  |  |
| Model resolution (Å) | 3.7 |  |  |
| FSC threshold | 0.143 |  |  |
| Model resolution range (Å) | 3.1-14.4 |  |  |
| Map sharpening *B* factor (Å²) | - 60 |  |  |
| Model-map correlation (box) | 0.74 |  |  |
| Model composition |  |  |  |
| Non-hydrogen atoms | 29832 |  |  |
| Protein residues | 1900 |  |  |
| Ligands | SY8: 1<br>ZN: 8<br>FE: 1 |  |  |
| *B* factors (Å²) |  |  |  |
| Protein | 75.72 |  |  |
| Ligand | 104.03 |  |  |
| R.m.s. deviations |  |  |  |
| Bond lengths (Å) | 0.003 |  |  |
| Bond angles (°) | 0.564 |  |  |
| Validation |  |  |  |
| MolProbity score | 1.74 |  |  |
| Clashscore | 4.99 |  |  |
| Poor rotamers (%) | 1.36 |  |  |
| Ramachandran plot |  |  |  |
| Favored (%) | 94.53 |  |  |
| Allowed (%) | 5.42 |  |  |
| Disallowed (%) | 0.05 |  |  |

**Figure 5.**
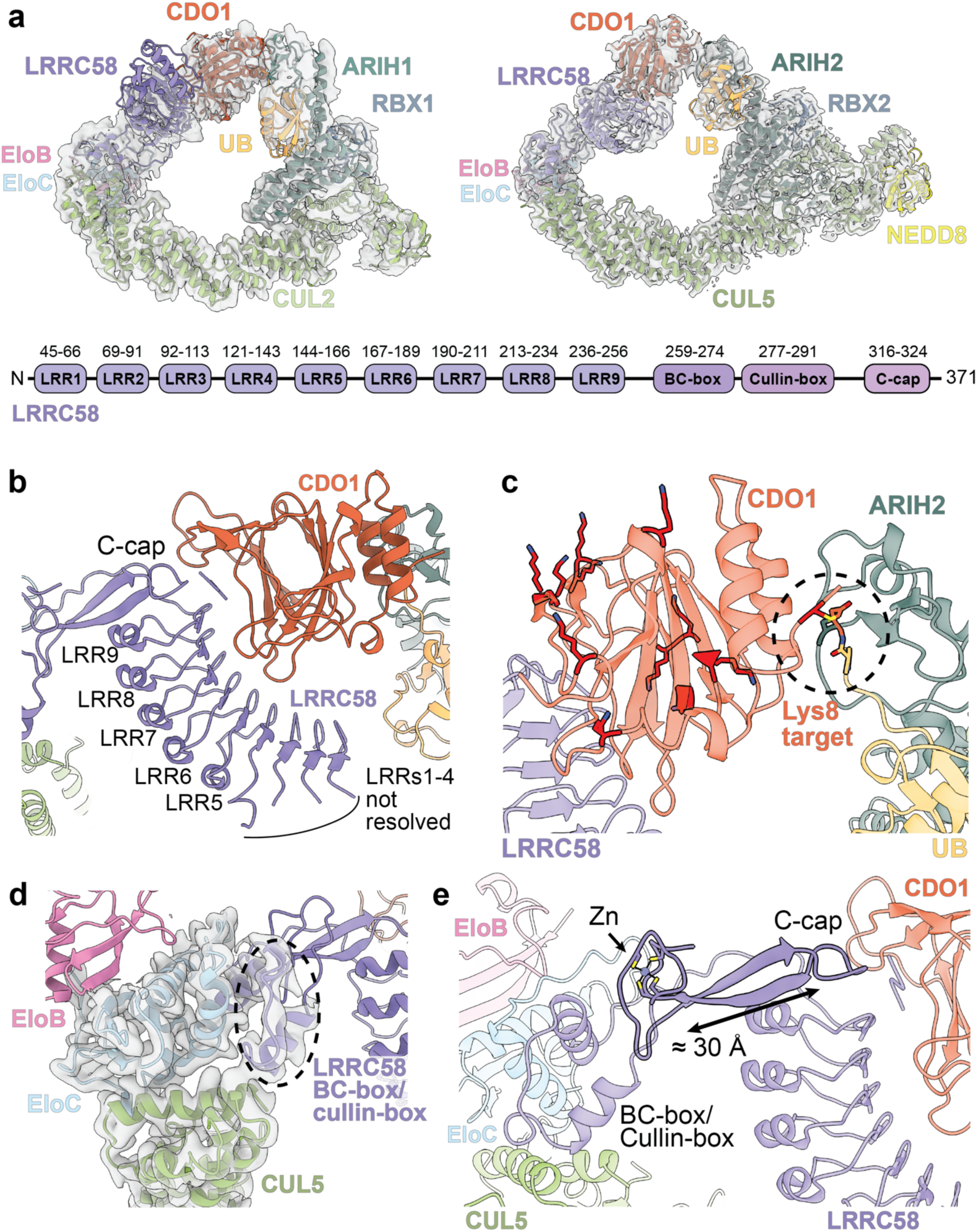
Cryo-EM structure of CDO1 engagement by LRRC58-CRL explains selective substrate Lys targeting. (a) The left panel displays a model of the sample representing CDO1 ubiquitylation by LRRC58 with neddylated CUL2-RBX1-ARIH1. Structural coordinates from EloB/C-CUL2 N-terminal domain (pink, light blue, and light green, respectively; PDB: 8WQH), CUL2 C-terminal domain (light green; PDB: 8Q7R), N-terminal ARIH and RBX1 (slate blue; PDB: 7B5M), and from AlphaFold3 models of the CDO1-LRRC58 complex (orange and purple, respectively) and ARIH1∼Ub (dark teal and gold, respectively) were fit into the corresponding composite EM density (transparent grey). The right panel displays the structure representing CDO1 ubiquitylation by LRRC58 with neddylated CUL5-RBX2-ARIH2 (EloB in pink, EloC in light blue, LRRC58 in purple, CDO1 in orange, NEDD8 in yellow, CUL5 in light green, Ub in gold, RBX2 in slate blue, ARIH2 in dark teal) and the corresponding composite EM density (transparent grey). Electron density for NEDD8 is not apparent in the CUL2 complex, as consistent with previous structures ^43,44^ due to the differing role of NEDD8 in CUL5-RBX2 versus CUL2-RBX1. Shown below is the schematic of domains from LRRC58 highlighting the LRR domains, BC-and Cullin-boxes, and the C-cap region. (d) Close up of the extensive interface between LRRC58 (purple) and CDO1 (orange) in the neddylated CRL5 complex. (e) Close-up of CDO1 (transparent orange) engagement of the catalytic Rcat domain of ARIH2 (LRRC58 in transparent purple, ARIH2 in transparent dark teal, Ub in transparent gold). CDO1 Lys side-chains are shown as opaque sticks. The Lys 8 target position, here replaced by a cysteine-substitution to generate a stable mimic of the ubiquitylation intermediate to facilitate cryo-EM structure determination, is highlighted (black dashed circle). (f) Close-up of the EloC-LRRC58-CUL5 interface with cryo-EM density (transparent grey) for the EloC and N-terminal CUL5 subunits (EloB in light blue, EloC in pink, CUL5 in light green, and LRRC58 in purple). (g) Close-up of LRRC58’s 2-stranded β-sheet (∼ 30 Angstroms; opaque purple) from which emanate the cullin-binding regions (BC-and cullin-box), the Zn binding loop (opaque purple, side chains of Zn-coordinating cysteine residues shown), and the CDO1 binding region (C-cap highlighted in opaque purple) (CUL5 in transparent light green, EloB in transparent light blue, EloC in transparent pink, CDO1 in transparent orange).

The superior map quality allowed building atomic coordinates for the sample representing CDO1 ubiquitylation by LRRC58 with neddylated CUL5-RBX2-ARIH2 (Extended Data Fig. 7). This, together with previous structures, allowed us to dock subcomplexes to also generate a model representing ubiquitylation by CUL2-RBX1-ARIH1. Despite the lower map quality of the CUL2 complex, these models allow for structural comparisons to be made between two ubiquitylation machineries employed by LRRC58.

The cryo-EM reconstructions visualize how LRRC58 selectively mediates CDO1 targeting. First, in complexes with CUL2 and CUL5, LRRC58 participates in the canonical CRL architecture (Fig. 5a, Extended Data Fig. 8). Second, LRRC58 makes extensive interactions with CDO1 (Fig. 5a,b). Third, LRRC58 projects CDO1 towards the catalytic portion of a neddylated CRL (Fig. 5a,c). Fourth, ubiquitylation site selectivity results from the constellation of CDO1 residues facing a ubiquitylation active site. Lys8 is the only accessible lysine within 25 Å of the active site in either ubiquitylation assembly (Fig. 5c).

The structure of LRRC58 had not previously been experimentally determined. Our data show the N-terminal portion of LRRC58 forms nine LRRs which are capped by a unique domain at the C-terminus. This “C-cap” and several LRRs together form the substrate-binding domain (Fig.5b). The intervening BC- and cullin-box region binds EloB/C and a cullin (Fig. 5d).

Notably, the 3-way interface between EloC, the LRRC58 cullin-box, and cullin were well resolved in the map with CUL5 (Extended Data Fig 7c). Despite use of this cullin in the complex, comparing with other structures showed LRRC58 displays a canonical CUL2-box, closely resembling that previously determined for the complex between the BC-box protein FEM1B and CUL2 (Extended Data Fig. 9a)^70^. LRRC58’s CUL2-box may explain the preferential requirement for CUL2 in determining stability of the CDO1 reporter. Nonetheless, our data also show a CUL2-box structure is fully compatible with engaging CUL5.

The sequences flanking both sides of LRRC58’s cullin-binding region adopted a unique multifaceted and multifunctional structure, centered around a twisted, ≈30 Å long two-stranded β-sheet (Fig.5e). Loops emanating from one end of the sheet form a zinc-bound structure that connects to the cullin-binding region. At the other end of this β-sheet, both strands, the loop between them, and the C-terminus serve as the cap of the LRRs and as a portion of the CDO1-binding site, hence our naming this the C-cap (Fig. 5b,e). Despite the key role of the LRRC58 C-terminus noted in a parallel study^52^, we were unable to fully resolve this entire region in our maps. However, additional density was observed, resembling a molecular glue between LRR9, the C-cap, and CDO1. We placed the four C-terminal LRRC58 residues as an extended region in this density because AlphaFold3predicts these residues fit between LRR9 and the C-cap albeit in a different arrangement (Extended Data Fig. 9b)^71^.

### Structural basis for CDO1 ubiquitylation by LRRC58 CRLs

The “top side” of CDO1^72^ engages a continuous surface in LRRC58 formed by the C-terminal seven LRRs and the C-cap. These interactions primarily map to four patches on CDO1 (Fig. 6a). Patch-1 consists of a CDO1 loop displaying Asp168, Gln169, Arg170, and His173. Patch-1 contacts the LRRC58 C-cap centered around Tyr322. Patch-2, is located at the center of the top side of CDO1. Here, Arg141 and Glu143 and the backbone of the adjacent Gly82 contact LRRC58’s C-terminal LRR strand (specifically Arg243) and C-terminus. Patch-3 is dominated by the CDO1 loop containing His147, which traverses the fifth through seventh LRR from LRRC58. This His147 side-chain specifically inserts between numerous LRRC58 residues, and contacts Tyr172. Patch-4 - the CDO1 loop containing Gln55 and Tyr56 – binds the opposite side of the seventh through ninth LRRC58 LRRs. In addition, we term a fifth CDO1 region (centered around Gln99 and Glu149) the D-patch based on prior studies showing these residues at the heart of the degrader-induced interaction with VHL (Fig. 6b,c)^55^.

**Figure 6.**
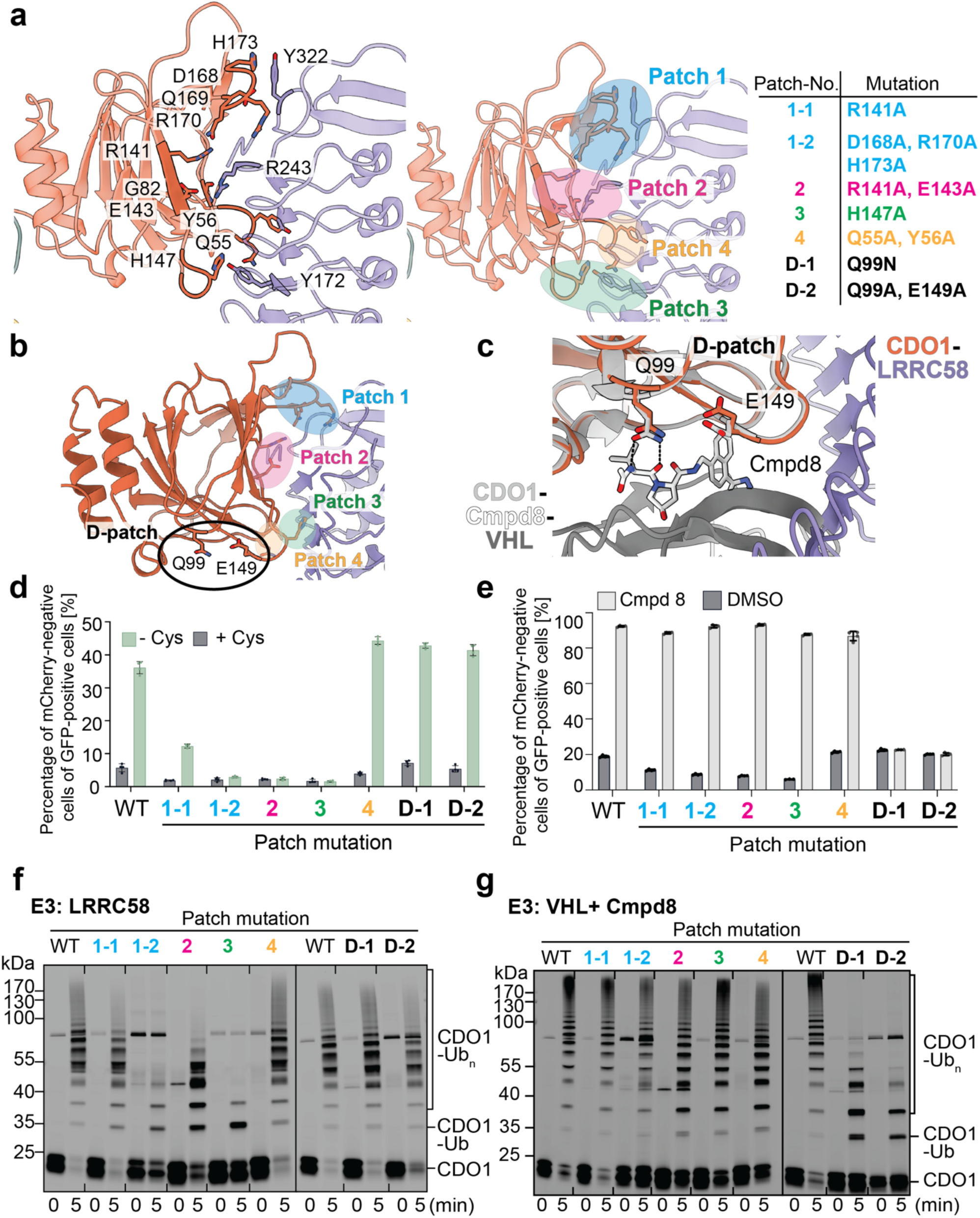
Perturbation of the LRRC58-CDO1 interface decreases CDO1 degradation in cells and in vitro ubiquitylation efficiency. (a) Intermolecular interactions along the LRRC58-CDO1 interface of the structure representing CDO1 ubiquitylation by LRRC58 with neddylated CUL5-RBX2-ARIH2. Residues involved from LRRC58 (purple) and CDO1 (orange) are labeled and shown in full opacity (right). The LRRC58-CDO1 interface has been sub-divided into patches as shown. Mutations made to these patches, and the D-patch (black) in panel (b) are summarized in the table at right. (b) Same as in (a) but highlighting the location of the D-patch residues, involved in degrader-induced interaction with VHL, on CDO1 with respect to the LRRC58 interface patches. (c) CDO1-Cmpd8-VHL-EloB/C crystal structure (PDB: 8VL9, CDO1 in light grey, VHL in dark grey, Cmpd8 in white) aligned to the CDO1 (orange) of the sample representing CDO1 ubiquitylation by LRRC58 (purple) with neddylated CUL5-RBX2-ARIH2. Side chains of D-patch residues Q99 and E149 are shown in sticks with hydrogen bonds between Cmpd8 and the CDO1-VHL residues in black dotted lines. (d) Reporter assay showing CDO1 variants expressed in HEK293T cells grown in complete or cysteine-free media. Only CDO1 with patch 1-3 mutations lose sensitivity to cysteine starvation whereas patch 4 or D-patch mutations respond to cysteine starvation. Average values from n=4 independent replicates are shown. Error bars report the standard deviation of the data points. (e) Reporter assay same as in (d) except comparing cells gown in complete media and treated with Compound 8 (Cmpd8) or DMSO. All CDO1 variants are efficiently degraded, except for those with mutations in the D-patch. Average values from n=4 independent replicates are shown. Error bars report the standard deviation of the data points. (f) In vitro reconstituted assays comparing WT and mutant Cy5-labeled CDO1 ubiquitylation in the presence of neddylated LRRC58-CUL2. Mutations in patch interfaces 1, 2, and 3 – but not in patch 4 or the D-patch – reduced CDO1 ubiquitylation. (g) Same as in (f) except with neddylated VHL-CUL2 and with DMSO or Cmpd8. CDO1 patch mutants show WT-like ubiquitylation efficiencies but the Cmpd8 binding site mutants display reduced ubiquitylation compared to WT.

Roles of the structurally-observed interactions were tested by examining effects of Ala substitutions in each CDO1 patch using our stability reporter system. Mutation of patches 1, 2, and 3 strongly impaired CDO1 instability induced by cysteine-starvation, but not that triggered by the VHL-dependent compound-8 (Fig. 6d,e). Furthermore, during preparation of our manuscript, unbiased saturation mutagenesis in a study posted on bioRxiv independently validated the roles of all four patches on cysteine starvation-induced degradation of similar CDO1 stability reporters^52^. This study also validated the roles for the LRRC58 residues binding CDO1 patches 1-3. Additionally, CDO1 patch-3 was also shown to be important for LRRC58-dependent degradation^54^.

We uniquely queried the mutants for effects on compound-8-induced degradation. These experiments served both as quality control for the stability reporters, and tested specificity. Our data showed these mutational defects were specific for the metabolically-signaled degradation pathway. Conversely, the D-patch variants were refractory to destabilization induced by compound-8, but were unstable under cysteine starvation conditions (Fig. 6d,e).

To gain mechanistic insights into the differential targeting of CDO1, we used purified proteins to compare effects of CDO1 mutations on ubiquitylation by the LRRC58- and compound-8/VHL-CUL2 CRLs (Fig. 6f,g). Importantly, the mutational effects on the two distinct ubiquitylation pathways paralleled those on the two distinct cellular degradation pathways.

Finally, the pair of structures provided insights into the ubiquitylation of a single substrate recruited by a receptor that can employ different cullins. We compared the showing CDO1 ubiquitylation by LRRC58-C with neddylated CUL2-RBX1-ARIH1 with the prior structure representing substrate ubiquitylation by a neddylated CUL1-RBX1-ARIH1 complex^44,73^ (Extended Data Fig. 8a). Remarkably, the catalytic ARIH1∼ubiquitin portion of the prior structure with CUL1 fit well in the map representing this assembly ubiquitylating CDO1. The same catalytic architecture was also observed in published maps for another CUL1 complex, and another map for a complex representing ubiquitylation by KHLDC10-CUL2 (Extended Data Fig. 8b,c). These data demonstrate that various neddylated RBX1-based CRLs can mediate substrate ubiquitylation by a generalizable ARIH1∼ubiquitin catalytic architecture.

Neddylated CUL5-RBX2-ARIH2 complexes also show a generalizable core assembly. Although there is no prior structure representing substrate ubiquitylation by neddylated CUL5-RBX2-ARIH2, there is precedent for such an assembly that did not have ubiquitin at the ARIH2 active site^74^. In the prior structure, the ARIH2 catalytic domain was not visible. Nonetheless, the neddylated CUL5-RBX2-ARIH2 that were visible superimpose with the LRRC58-CDO1 complex (Extended Data Fig. 8d).

Unexpectedly, however, aligning the cullins in the two LRRC58 complexes show different relative arrangements for the two key units underlying ubiquitylation. The LRRC58-CDO1 receptor-substrate complex is oriented differently compared to the N-termini of CUL2 and CUL5 (Extended Data Fig. 10a). The different placements of the substrate are accommodated by different orientations of the ARIH1 and ARIH2 active sites (Extended Data Figs. 8d,e,10a). Thus, these data suggest a distinctive catalytic arrangement for CUL5-ARIH2, although functional importance will depend on future studies.

## Discussion

Our data provide a molecular pathway for elegant regulation, discovered decades ago, through degradation of CDO1 by the ubiquitin-proteasome system when cysteine is limiting^35–37^. Now our proteomics-based strategy has identified co-regulation of a cullin-RING ligase receptor, LRRC58, and CDO1 as its substrate (Fig. 1,2). We found that cysteine starvation protects LRRC58 from ubiquitin-mediated degradation (Fig. 1,2). This in turn controls CDO1 through its direct ubiquitylation and degradation (Fig. 2).

Through cellular and biochemical reconstitution, we defined facets underlying CDO1 regulation by LRRC58 and by the recently developed molecular glue degrader compound-8 with VHL. While the native LRRC58-dependent pathway preferentially targets CDO1 Lys8 for ubiquitylation, the degrader molecule overrides this through ubiquitylation on multiple sites (Fig. 4). Cryo-EM data support that LRRC58 binds a different side of CDO1 in comparison with the VHL/compound-8 complex (Fig.6a,b). Importantly, LRRC58 projects CDO1 Lys8 towards the ubiquitylation active site. Although future structural studies are required to show how the VHL/compound-8-bound CDO1 undergoes ubiquitylation, its functional differences relative to the LRRC58 pathway highlight how distinct presentations of a single substrate to a common cullin can tune ubiquitylation and degradation efficiencies.

Distinctions in native versus drug-mediated substrate ubiquitylation is of relevance towards successful development of therapeutics mediating targeted protein degradation. For instance, we note that two of three CDO1 mutations recently reported as the only genetic etiology discovered in patients with a rare neurological disorder map to CDO1 patches 2 and 3 (Glu143 and His147) contacting LRRC58 (Fig. 6a)^75^. While the data here show variants in these residues are defective for LRRC58-dependent ubiquitylation under cysteine starvation, the same CDO1 mutants are targeted by compound-8/VHL(Fig. 6d-g). Thus, our data indicate that disease-associated proteins could be targeted by orthogonal recognition induced by degrader molecules.

Our structural data also revealed unexpected plasticity for LRRC58’s cullin-box, which can bind CUL5 as well as CUL2. In combination with previous work, the structures elucidated here define a consensus active configuration for ARIH1-mediated ubiquitylation of substrates recruited to neddylated CUL1 and CUL2 E3s, and a twist in ARIH2 for the corresponding ubiquitylation complex with substrate recruited to a neddylated CUL5 E3 (Extended Data Figs. 8d,e).

While our manuscript was in preparation, two other studies also reported human LRRC58 regulation of CDO1. The confluence of independent data supports the robustness of our findings, and highlights the range of methodologies available to discover metabolic signaling through the CRL pathways. One study relied on an elaborate approach - correlating levels of 285 metabolites with 11,868 proteins across genetically diverse mouse and human samples - to discover linkage between taurine and hypotaurine, LRRC58, and CDO1^54^. The other study employed ubiquitin-focused CRISPR screening. This preprint reported the role of LRRC58-CUL2 in destabilizing a fluorescent CDO1 stability reporter under cysteine-limiting conditions (and in destabilizing a stability reporter for LRRC58 itself in cysteine-rich conditions)^52^. The orthologous *C. elegans* pathway was also recently reported^53^. A series of genetic screens and epistasis experiments initially pursuing differences in *C. elegans* growth on various bacteria ultimately indicated that LRRC58 controls sulfur metabolism through post-translational regulation of CDO1.

By contrast, while our initial results relied on proteomic strategies we developed to reveal responses to signals^38^, our follow-up experiments showed that the relationship between levels of cysteine, LRRC58 and CDO1 were apparent by straightforward proteome analysis by DIA-MS combined with high-throughput computational modeling.

Retrospectively, our new pipeline is consistent with prior findings that many CRL substrate receptors are regulated by autodegradation that is suspended when these E3s are needed to ubiquitylated substrates^38,46,47,56–58^. Our study further highlights how proteomics can shed light on E3 targets, through comparison of total proteomes and modeling of protein complexes. This strategy can be universally applied across cell lines and perturbations, and it yields data on endogenous proteins without requiring sophisticated metabolomics, elaborate computation, or CRISPR screening to point towards new pathways.

Now, CDO1 joins IDO1 (indoleamine 2,3-dioxygenase 1) and TDO2 (tryptophan 2,3-dioxygenase) as oxidoreductase enzymes that are downregulated by a CRL in response to metabolic stress ^76,77^. IDO1 is destabilized when heme biosynthesis is inhibited. Recent studies showed how the level of IDO1 is controlled in a heme-dependent manner by the KLHDC3-CUL2 E3^77^. KLHDC3-CUL2 preferentially ubiquitylates a lysine near the C-degron of apo-IDO1. However, this IDO1 region is reshaped upon heme binding, allosterically hindering accessibility to the CRL. Meanwhile, the abundance of TDO2, like CDO1, is regulated by availability of its amino acid substrate^78^. Degradation of these enzymes when their respective tryptophan or cysteine substrate is limiting maintains homeostasis by preventing toxic depletion of a crucial amino acid. TDO2 is stabilized by its directly binding to tryptophan, at an “exosite” distal from the active site^79^. Under conditions of low tryptophan, TDO1 is subject to CUL1-dependent degradation. However, understanding how amino acid binding to the exosite prevents TDO2 degradation awaits a report of its substrate receptor.

A distinct mode of metabolite sensing is now suggested from the inhibition of LRRC58-mediated degradation of CDO1 when cysteine is plentiful. However, the cysteine sensing mechanism remains unclear. LRRC58 residues serving as a “cysteine sensor” were identified by saturation mutagenesis of an LRRC58 stability reporter described in a recent preprint^52^. However, mapping those residues onto our structure of the LRRC58-CDO1 complex did not reveal an amino acid-binding motif (Extended Data Fig. 10b). On the other hand, the structure shows that some “cysteine sensor” residues promote LRRC58 binding to CDO1, raising the possibility that substrate binding protects LRRC58 from degradation. Such regulation does not explain how cysteine levels are percieved at a molecular level. Other “cysteine sensor” residues map to a cysteine-rich Zn-binding element, which orients LRRC58’s substrate relative to the cullin and ubiquitylation active site (Extended Data Fig. 10b). These residues could potentially play a role in projecting LRRC58 for ubiquitylation, although deciphering the cysteine sensing mechanism will require further investigation. Irrespective of the molecular mechanism, the regulation of CDO1 through metabolic control of LRRC58 stability (Fig. 1,2) provides yet another means for regulation of an oxidoreductase enzyme by its amino acid substrate. It seems likely that widespread control of metabolic enzymes through the ubiquitin-proteasome system awaits discovery. We anticipate that crosstalk between the ubiquitin system and metabolism will be unearthed by a plethora of technologies – genetics^53^, multi-sample multiomics^54^, CRISPR screening^52^, and proteomics together with structural modeling (Fig. 1,2) – just as these strategies all converged to decipher the LRRC58 response to cysteine abundance.

## Acknowledgements

We thank Charlotte Duteil for advice on flow cytometry, Jakob Farnung for advice on assay design, Sara Šepić and Rajan Prabu for advice on cryo-EM processing and Josef Kellermann and all other members of the Schulman lab for general advice and assistance.

We also thank Barbara Steigenberger and Victoria Sánchez at the MPIB Mass Spectrometry Facility (RRID:SCR_025745), Rinho Kim at the MPIB NGS Core Facility (RRID:SCR_025746), Assa Yeroslaviz at the Bioinformatics Core Facility (RRID:SCR_025742), Martin Spitaler and Markus Oster at the MPIB Imaging Facility (RRID:SCR_025739), Stephan Uebel and Stefan Pettera at the MPIB Biochemistry Core Facility (RRID:SCR_025743) and Daniel Bollschweiler and Tillman Schäfer at the MPIB Cryo-EM Facility (RRID:SCR_025744).

This study was co-funded by the Max Planck Society and the European Union (ERC, UPSmeetMet, 101098161, BAS). LS, AT, and SAM were supported by PhD fellowships from the Boehringer Ingelheim Fonds. Views and opinions expressed are however those of the authors only and do not necessarily reflect those of the European Union or the European Research Council. Neither the European Union nor the granting authority can be held responsible for them.

## Author contributions

Biochemistry: G.A.A, L.S; Cryo-EM, structure building and refinement: G.A.A., L.S.; Proteomics: G.A.A., L.T.H; Cell biology: K.S., A.T., P.J.M., G.A.A., C.G., J.M.; Protein production: L.S., G.A.A, L. Sc, S.A.M, J.D, S.vG, C.S; Data analysis: G.A.A, L.S, K.S., A.T., L.T.H, B.A.S, P.J.M; Conception: P.J.M; Technical conception: A.T., L.T.H, B.A.S, G.A.A, L.S, K.S; Supervision: B.A.S, P.J.M, M.M, G.K; Paper preparation: G.A.A, L.S., G.K, P.J.M, B.A.S, with input from all authors

## Competing interests

BAS is a member of the scientific advisory boards of Proxygen and Lyterian. The other authors declare no competing interests.

---

**Extended Data Figure 1.**
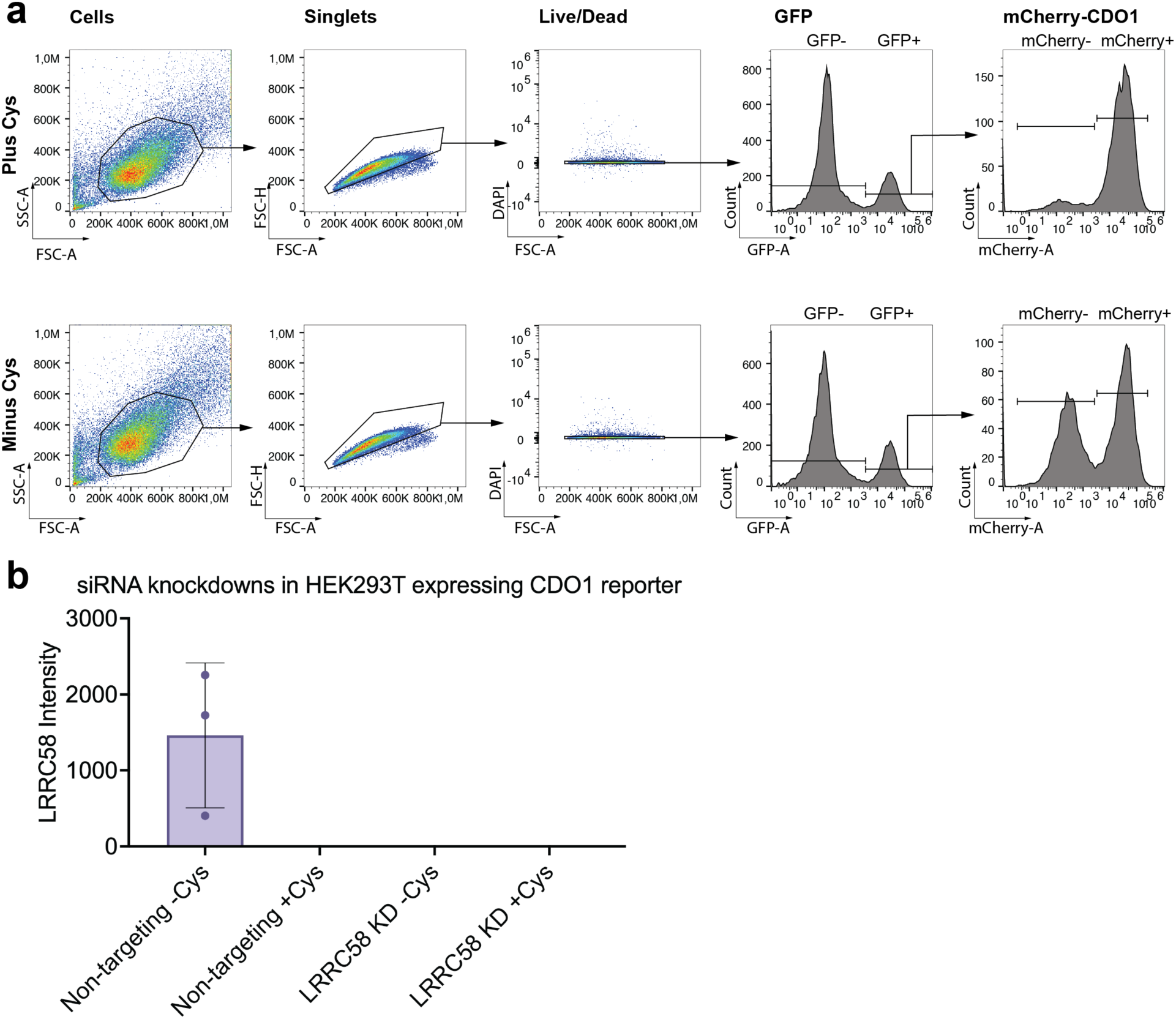
CDO1 stability reporter array. (a) Representative gating strategy for HEK293T CDO1 reporter cells analyzed in FlowJo. Cells were first gated using FSC-A/SSC-A to exclude debris, followed by doublet discrimination using FSC-A/FSC-H. Live cells were identified based on DAPI negativity. A histogram of the live-cell population was then used to gate GFP-positive (GFP⁺) reporter cells, within which mCherry-positive (mCherry⁺) and mCherry-negative (mCherry⁻) subpopulations were defined. Changes in mCherry-signal were subsequently used to assess CDO1 degradation. (b) LRRC58 knockdown (KD) efficiency was determined through analysis of the total proteome of HEK293T cells expressing CDO1 reporter in the absence and presence of extracellular cysteine (-/+ Cys), as compared to the non-targeting control.

**Extended Data Figure 2.**
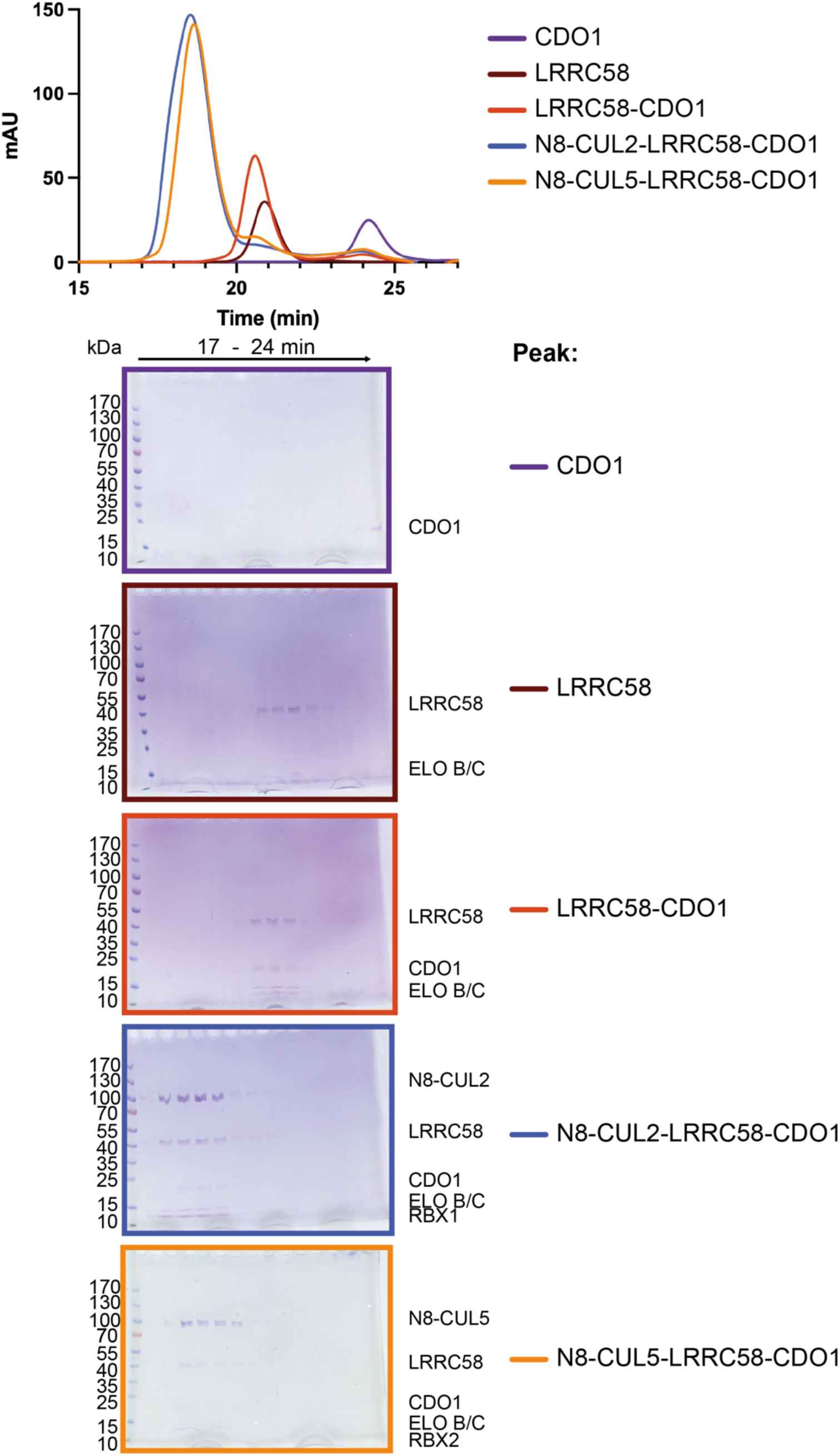
LRRC58-EloB/C-CDO1 complex stably interacts with neddylated CUL2-RBX1 and neddylated CUL5-RBX2. Graph showing UV absorbance versus Superdex 200 Increase 3.2/300 gel filtration column retention time for the indicated protein samples that had been mixed stoichiometrically (5 µM) prior to injection. Shifts in the peak retention time for substrate receptor-containing CRL2 and CRL5 complexes compared to traces for individual sub-components, indicating complex formation. Coomassie stained SDS-PAGE gels (bordered in colors corresponding to the chromatograms shown above) show the protein components from the indicated gel filtration fractions.

**Extended Data Figure 3.**
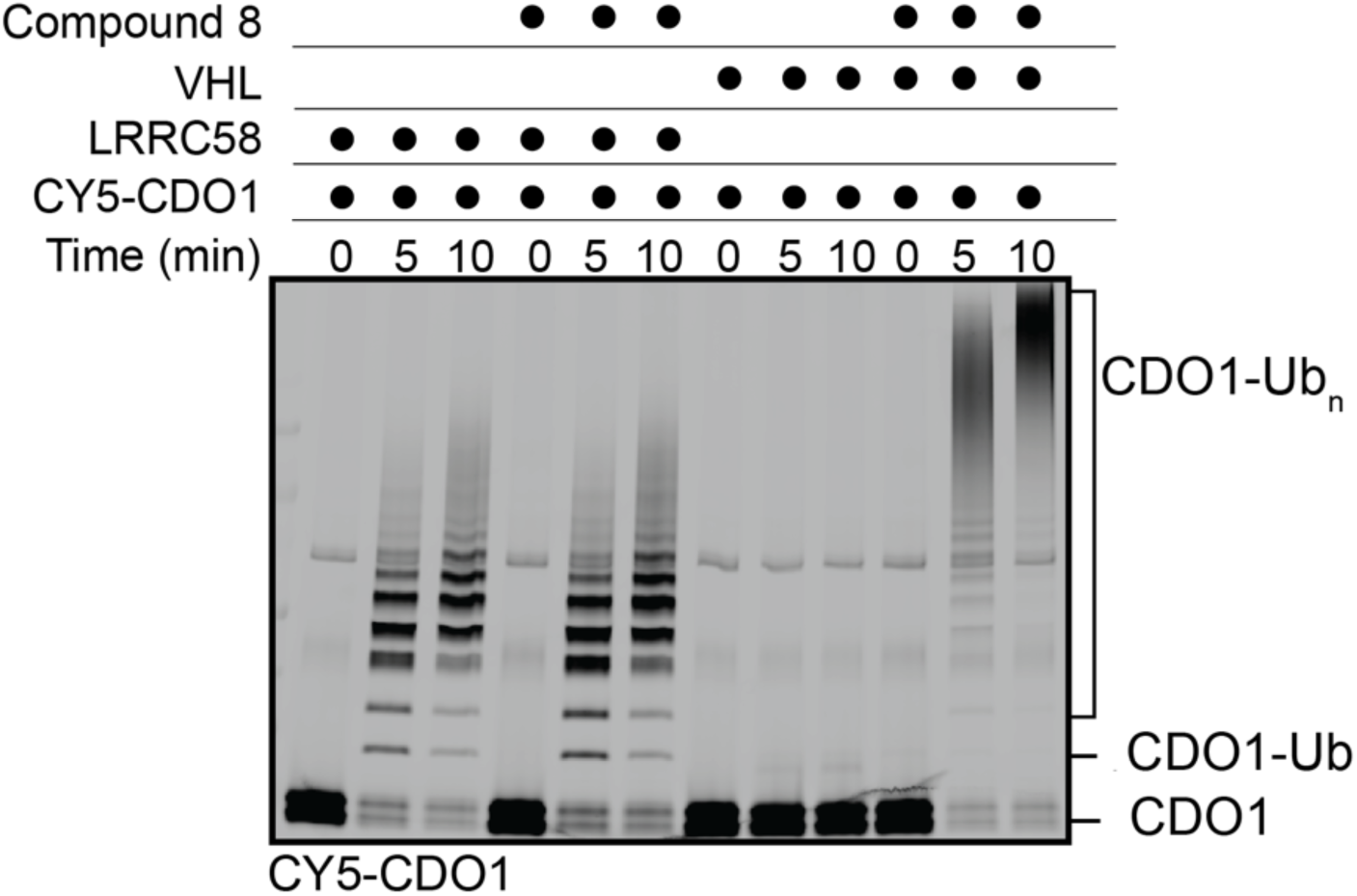
The molecular glue degrader, Compound 8, promotes CDO1 ubiquitylation to a greater extent than the native pathway. In vitro reconstituted ubiquitylation reactions containing Cy5-labeled CDO1 with neddylated CUL2-RBX1 and in the absence or presence of LRRC58-EloB/C (native pathway) or VHL-EloB/C (degrader pathway) and compound 8 or DMSO control. Compound 8 does not increase CDO1 ubiquitylation with LRRC58-EloB/C as substrate receptor but massively increases CDO1 ubiquitylation with VHL-EloB/C. Fluorescence scans are representative of n=3 technical replicates.

**Extended Data Figure 4.**
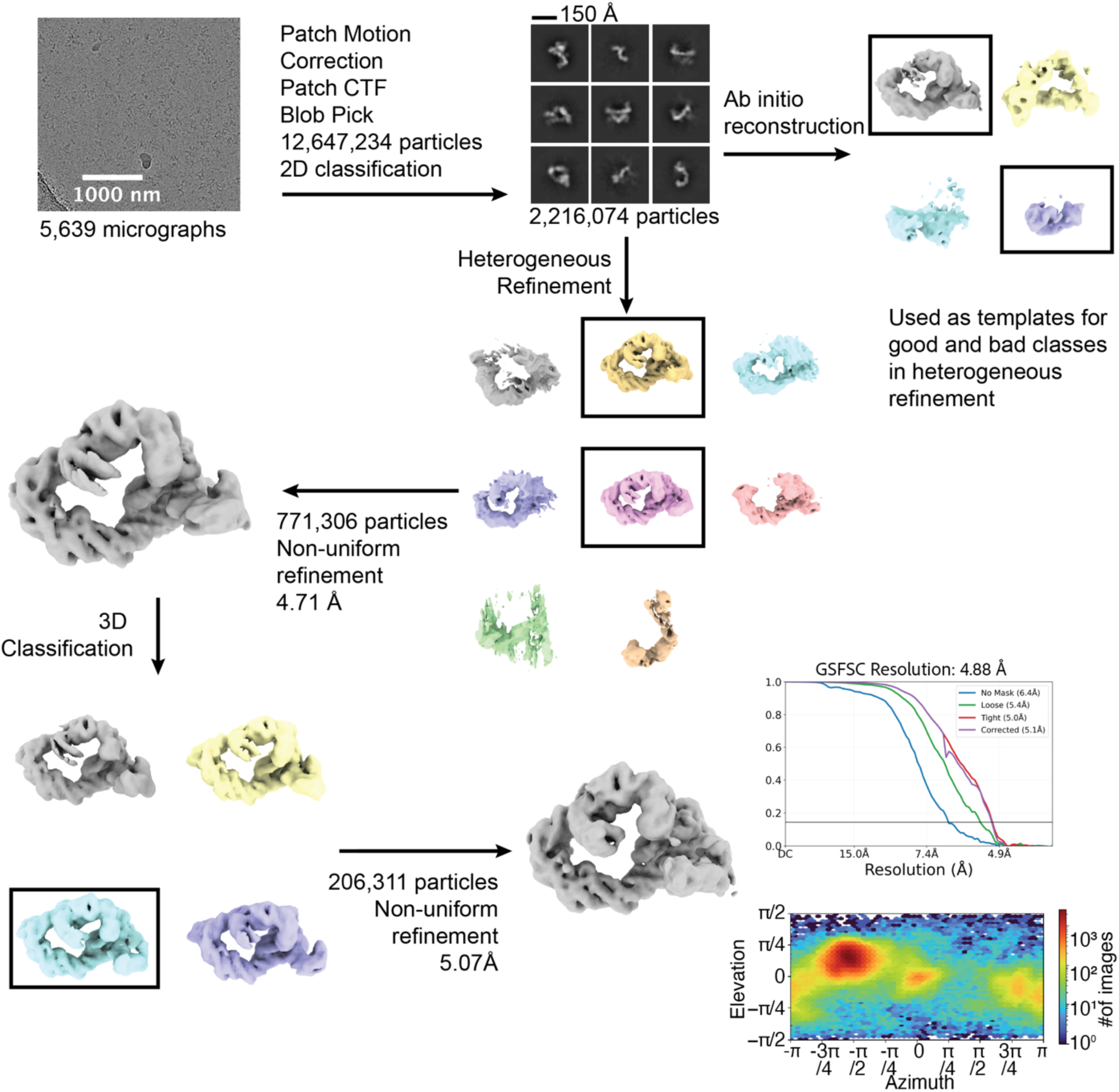
Cryo-EM processing scheme for screening dataset for sample representing CDO1 ubiquitylation by LRRC58 with neddylated CUL2-RBX1-ARIH1. Data processed in CryoSPARC v4.7.1 yielded a 3D reconstruction with a resolution of 5.07 Å (as determined by the gold-standard Fourier shell correlation of 0.143, shown). Orientation distribution plot of particles from final refinement also shown. This volume was used as input for Heterogeneous Refinement in Extended Data Fig 5.

**Extended Data Figure 5.**
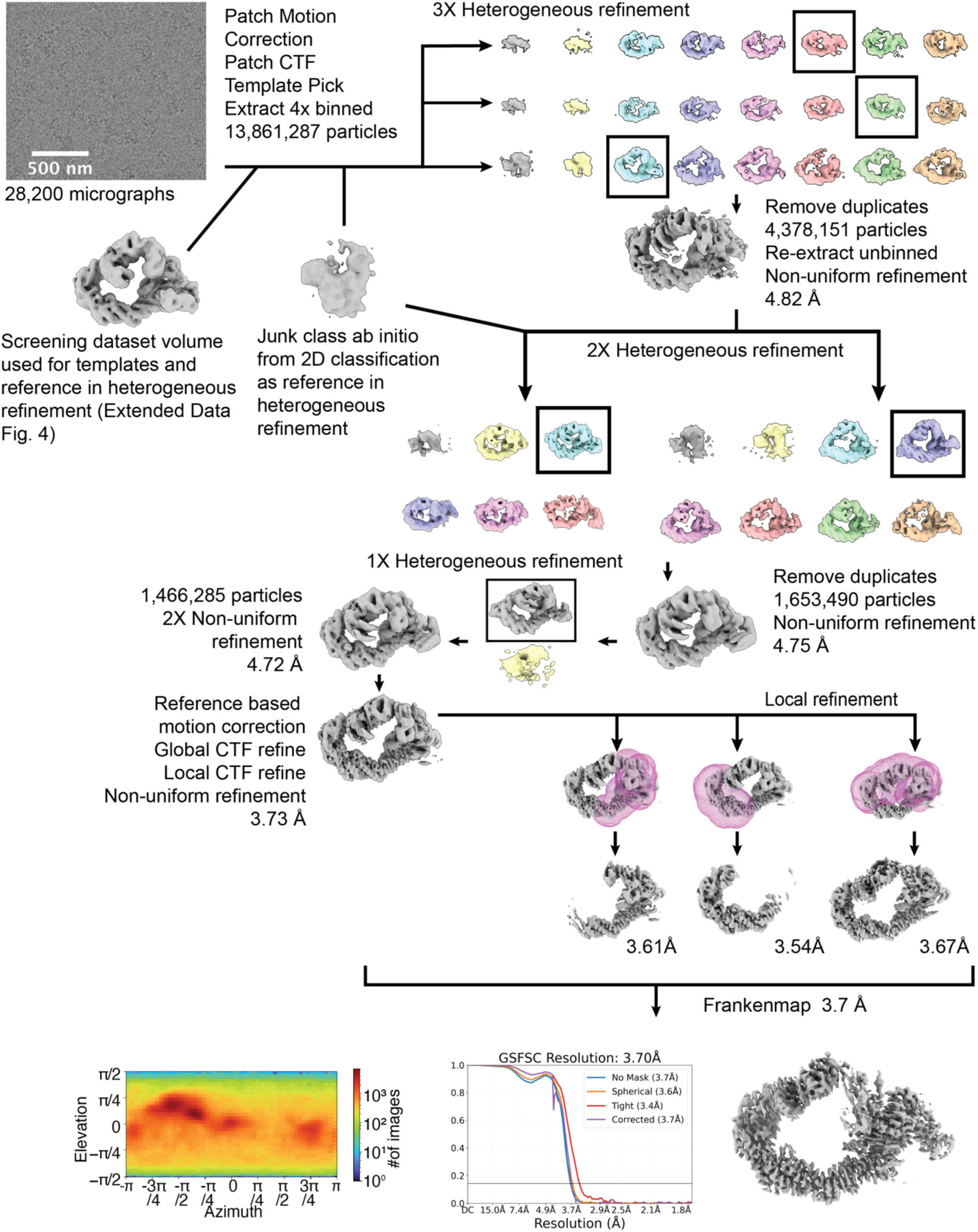
Cryo-EM processing scheme for sample representing CDO1 ubiquitylation by LRRC58 with neddylated CUL2-RBX1-ARIH1. Data processed in CryoSPARC v4.7.1 yielded a consensus 3D reconstruction with a resolution of 3.73 Å, and three locally refined maps with resolutions of 3.61 Å, 3.54 Å, and 3.67 Å. Locally refined maps were combined with the consensus map to create a 3.7 Å composite map (as determined by the gold-standard Fourier shell correlation of 0.143, shown) using Frankenmap (Warp v1.9.0). The orientation distribution plot of particles from consensus refinement is also shown.

**Extended Data Figure 6.**
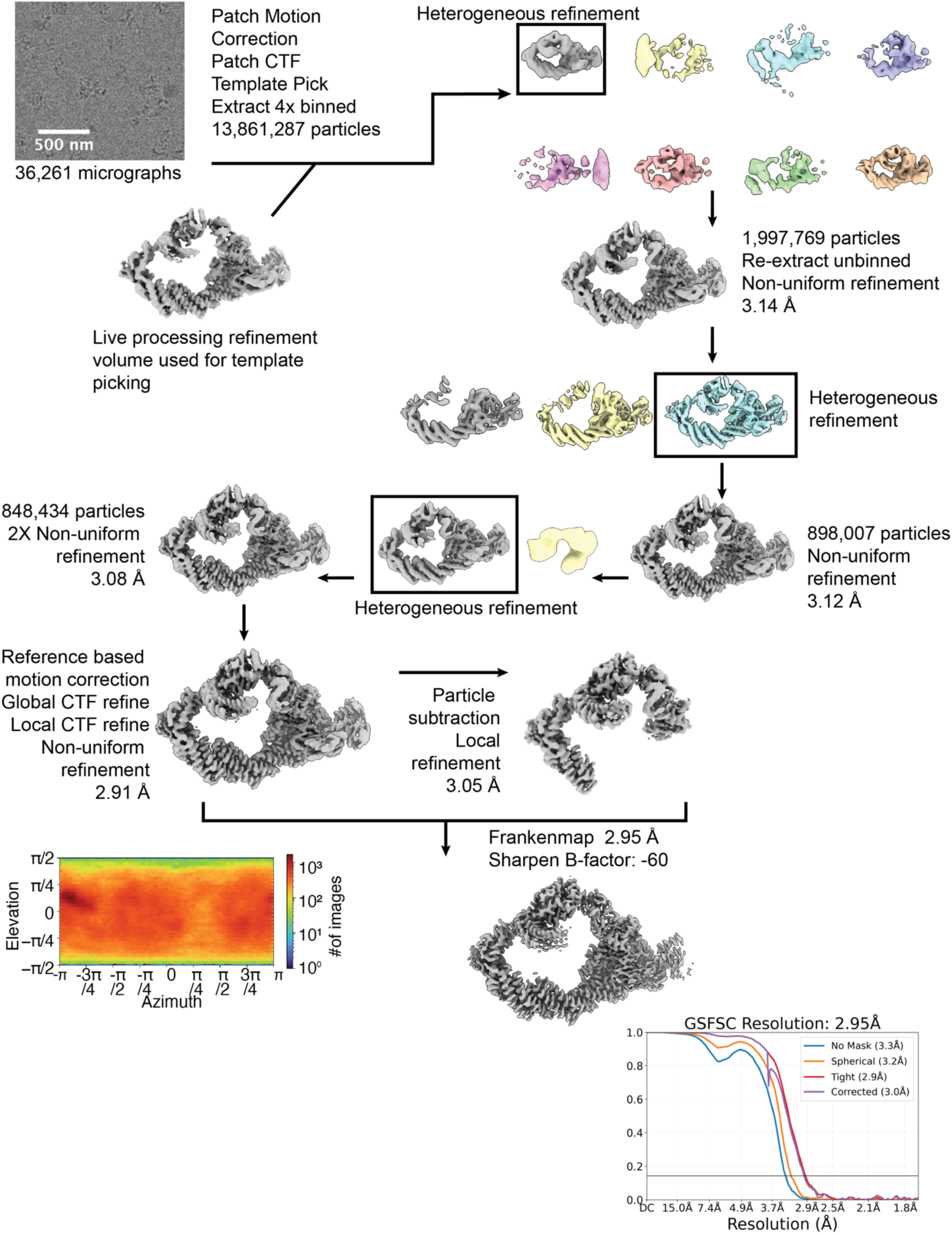
Cryo-EM processing scheme for sample representing - CDO1 ubiquitylation by LRRC58 with neddylated CUL5-RBX2-ARIH2. Data processed in CryoSPARC v4.7.1 yielded a consensus 3D reconstruction with a resolution of 2.91 Å, and a particle subtracted and locally refined map with a resolution of 3.05 Å. The particle subtracted map was combined with the consensus map to create a 2.95 Å composite map (as determined by the gold-standard Fourier shell correlation of 0.143, shown) using Frankenmap (Warp v1.9.0). The orientation distribution plot of particles from consensus refinement is also shown.

**Extended Data Figure 7.**
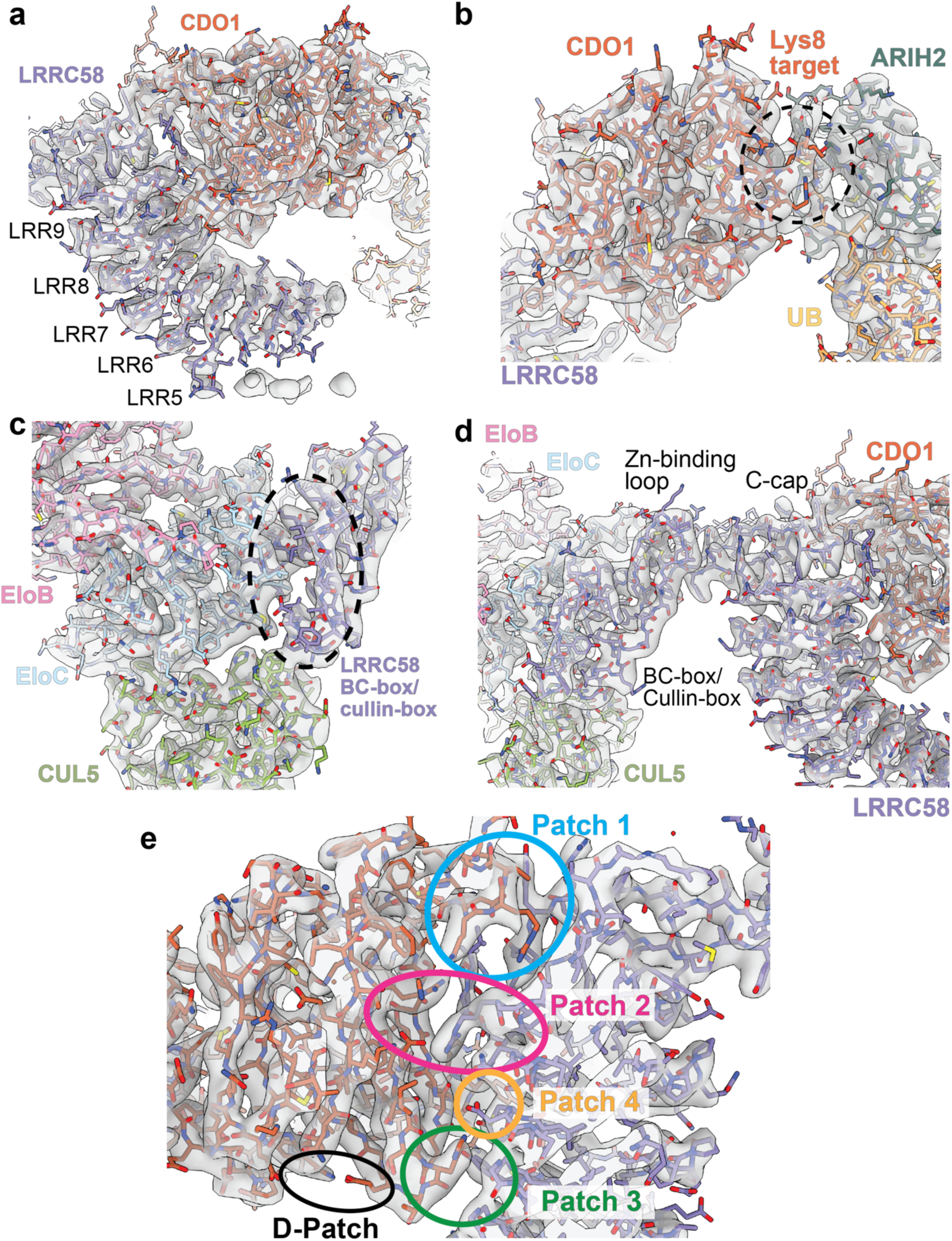
Example fits of the structure representing CDO1 ubiquitylation by LRRC58 with neddylated CUL5-RBX2-ARIH2 into the cryo-EM reconstruction. Atomic model representing the ubiquitylation intermediate in the CDO1-LRRC58 complex with CUL5-RBX2-ARIH2 shown in stick representation and fit in the corresponding composite cryo-EM map (transparent grey). (a) Overview of LRRC58 (purple) interaction with CDO1 (orange), highlighting the positions of the LRR regions that had sufficient electron density to enable their modeling. (b) Zoomed view of CDO1 (orange) highlighting the ARIH2 Rcat domain (LRRC58 in purple, ARIH2 in dark teal, Ub in gold). CDO1 Lys 8 target position, here replaced by a Cys-substitution to enable three-way cross-linking between ARIH2, ubiquitin and CDO1 (dashed black circle), mimicking the high energy ubiquitylation transition state intermediate. (c) Molecular details of the interfaces mediated EloC-LRRC58-CUL5 interaction (EloB in light blue, EloC in pink, CUL5 in light green, and LRRC58 in purple). LRRC58’s BC- and Cullin-boxes are identified by the dashed circle. (d) Same as (c) but with an expanded view including CDO1 (orange) and LRRC58’s C-cap and Zn binding loop regions. (e) Zoomed view highlighting the LRRC58-CDO1 intermolecular interface and various subregions (patch 1-blue, patch 2-pink, patch 3-green, patch 4-yellow, and D patch).

**Extended Data Figure 8.**
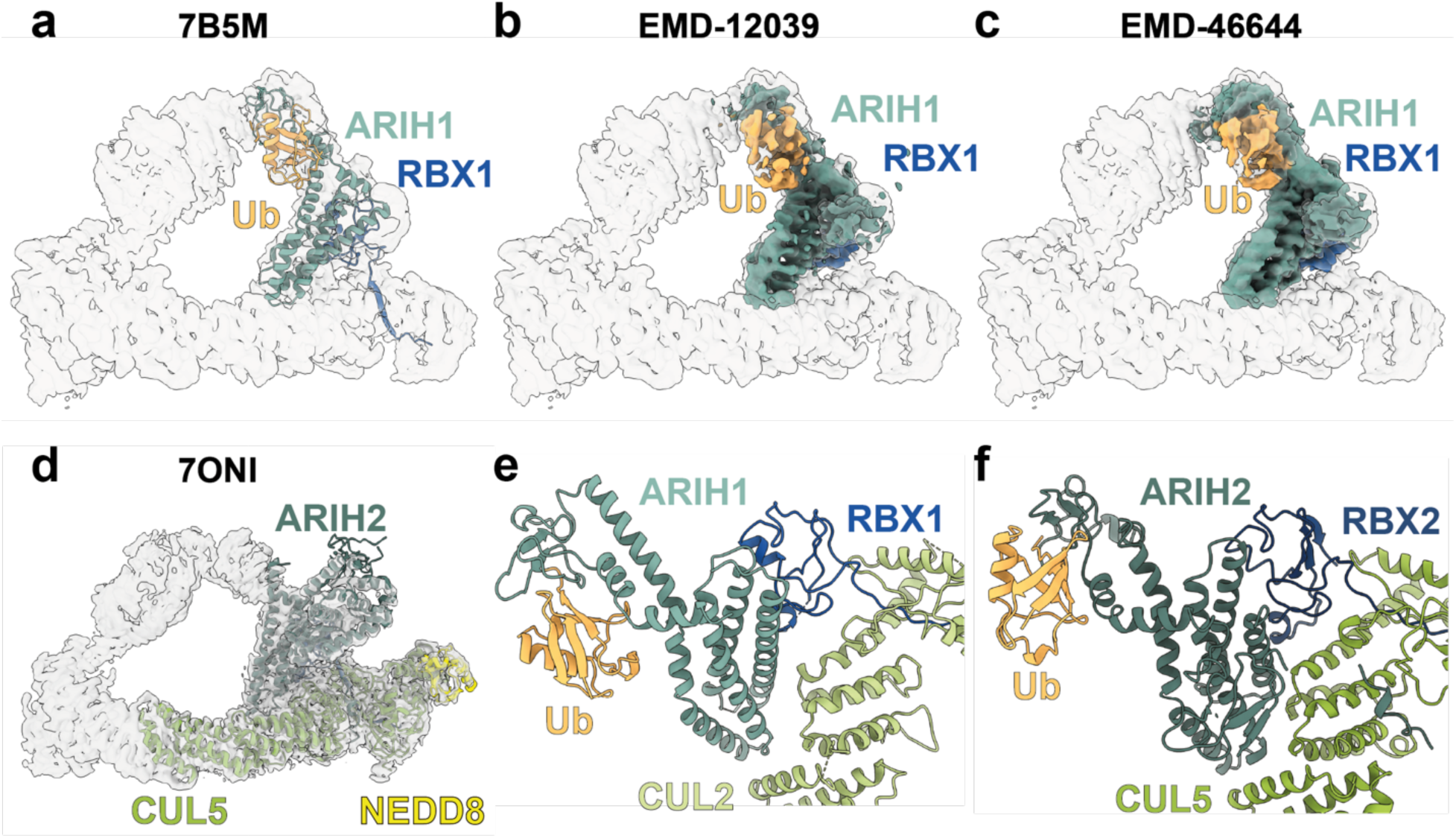
RBX1-based CRLs shows distinct catalytic arrangement from RBX2-based CRLs. (a) The RBX1-ARIH1∼ubiquitin portion of a CUL1 CRL structure (RBX1 in blue, ubiquitin in gold, and ARIH1 in green, PDB: 7B5M) fit into the composite map representing CDO1 ubiquitylation by LRRC58 with neddylated CUL2-RBX1-ARIH1 (transparent grey). (b) Same as in (a), but the RBX1-ARIH1∼ubiquitin region of a CUL1 CRL cryo-EM map (EMD-12039). (c) Same as in (a) and (b), but the RBX1-ARIH1∼ubiquitin region of a CUL2 CRL cryo-EM map (EMD-46644). (d) The CUL5 C-terminal domain (light green), ARIH2 (dark green), and NEDD8 (yellow) which did not have ubiquitin at the ARIH2 active site (PDB: 7ONI) fit into the composite map representing CDO1 ubiquitylation by LRRC58 with neddylated CUL5-RBX2-ARIH2 (transparent grey). (e) The RBX1-ARIH1∼ubiquitin portion of the model representing CDO1 ubiquitylation by LRRC58 with neddylated CUL2-RBX1-ARIH1, aligned on the ARIH2 domain of the RBX2-ARIH2∼ubiquitin portion in the structure representing CDO1 ubiquitylation by LRRC58 with neddylated CUL5-RBX2-ARIH2 in panel (f). The catalytic region is oriented differently between the two CRL complexes.

**Extended Data Figure 9.**
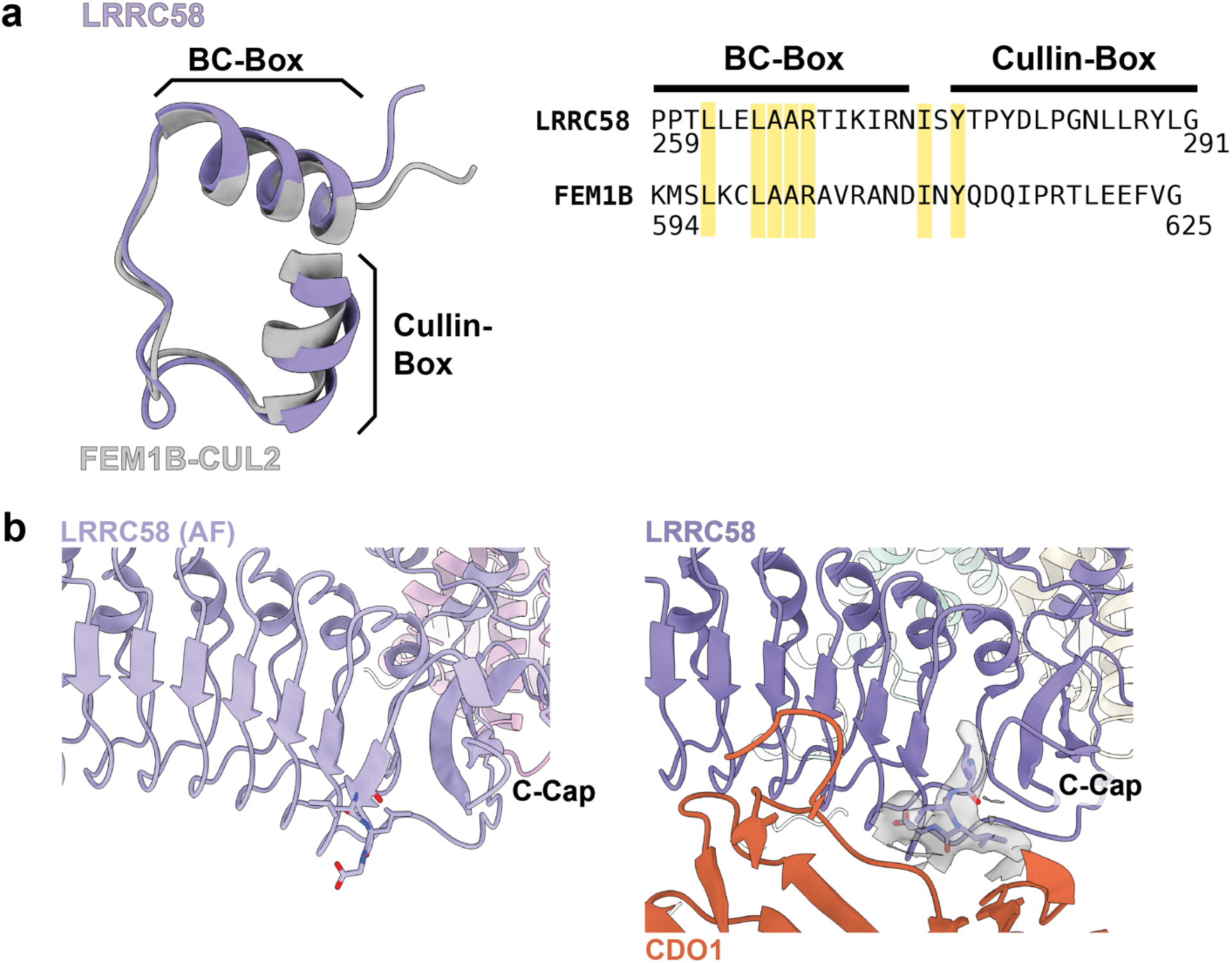
CUL2/5-dependent substrate receptors share conserved BC-box structures. (a) Ribbon diagrams showing structural alignment of the BC-and Cullin-Boxes from the LRRC58 with CUL2-dependent substrate receptor FEM1B’s BC-and Cullin-Boxes (grey, PDB: 8WQH). Amino acid sequences occupying structural equivalent positions from LRRC58 and FEM1C are aligned (identical residues are highlighted in yellow). (b) Displayed on the left, AlphaFold3 (AF) model of full-length LRRC58 (light purple, the last four residues of the C-terminal tail are displayed as sticks). For comparison, the LRRC58 (purple) -CDO1 (orange) interface is shown (right) in the structure representing CDO1 ubiquitylation by LRRC58 with neddylated CUL5-RBX2-ARIH2. While electron density for the full LRRC58 C-terminus was largely missing, sufficient density was noted at the extreme C-terminus (the composite map, shown in transparent grey) and enabled modeling of the final 4 residues of LRRC58’s C-terminal tail which showed similarity to the AF model.

**Extended Data Figure 10.**
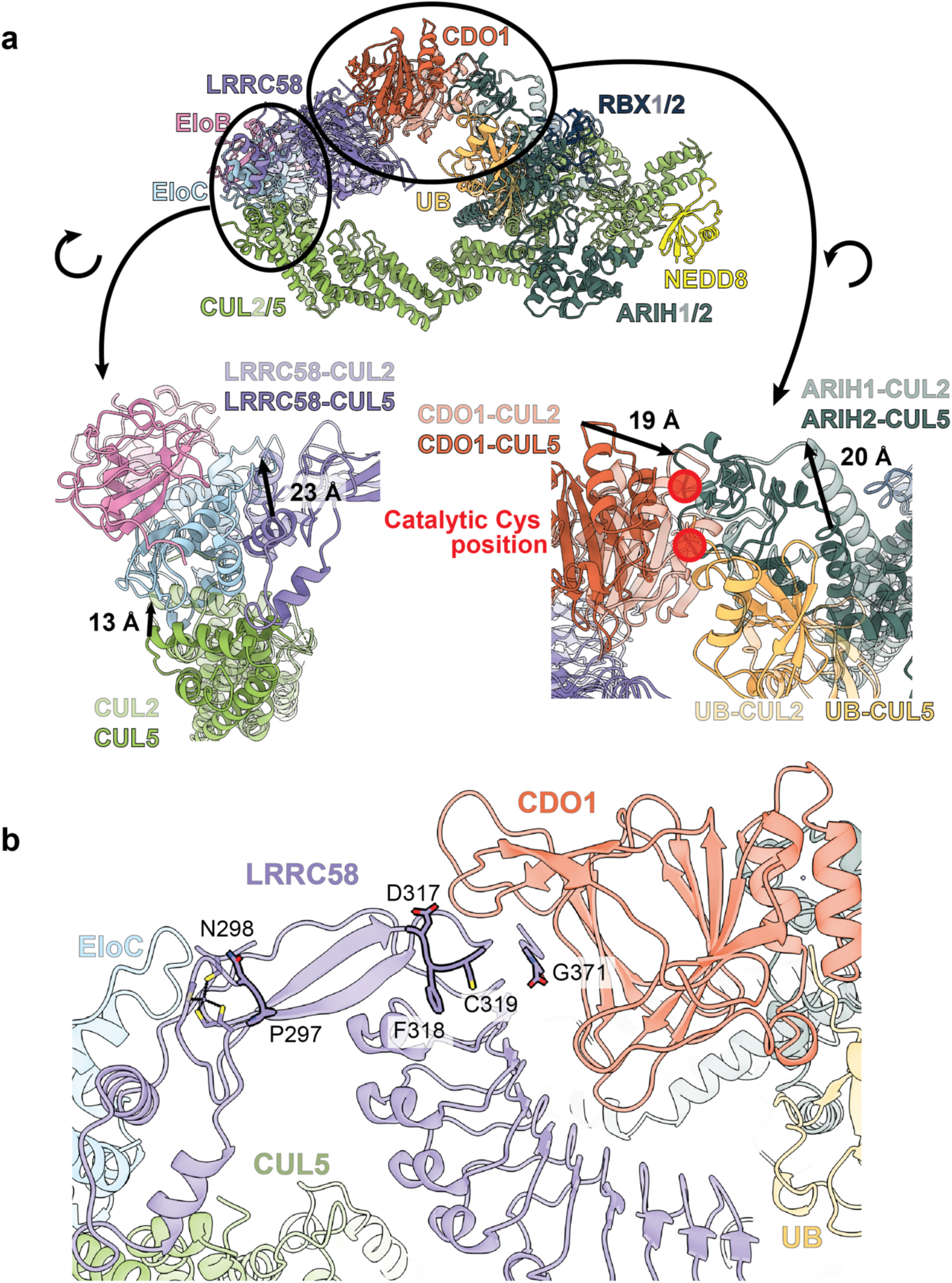
Conformational flexibility of LRRC58-CDO1 in complex with either neddylated CUL2 or CUL5 and their respective ubiquitin-carrying enzymes. (a) Ribbon diagram of the structure representing CDO1 ubiquitylation by LRRC58 with neddylated CUL5-RBX2-ARIH2 (EloB in pink, EloC in light blue, LRRC58 in purple, CDO1 in orange, NEDD8 in yellow, CUL5 in light green, Ub in gold, RBX2 in slate blue, ARIH2 in dark teal) aligned to the model representing structure representing CDO1 ubiquitylation by LRRC58 with neddylated CUL2-RBX21-ARIH1 (displayed in transparent ribbons, same colors as previous but with RBX1 in transparent blue and ARIH1 in transparent teal). Detailed comparisons are shown below the overall structural alignment, highlighting regions that displayed large magnitude structural shifts. CUL2 shifts up to 13 Å while the LRRC58 BC-box shifts up to 23 Å. In the catalytic region (catalytic cysteine residues (357 for ARIH1, 310 ARIH2) highlighted with red circles), CDO1 shifts upwards of 19 Å and ARIH1 shifts 20 Å from the ARIH2 position in the CUL5 complex. (b) Mapping of LRRC58 residues (in opaque sticks, outlined in black), identified by saturation mutagenesis by Ramage et al., 2025, that may function as cysteine sensors. Residues are shown on the structure representing CDO1 ubiquitylation by LRRC58 with neddylated CUL5-RBX2-ARIH2 (in transparent ribbons; LRRC58 in purple, EloC in light blue, CDO1 in orange, CUL5 in light green, Ub in gold, ARIH2 in dark teal). The “cysteine sensor” residues either interact with CDO1 or are located close to the LRRC58 Zn-binding loop. Residues L330 and M365 were identified but are not visualized in our structure.

## Notes

### Competing Interest Statement

B.A.S. is a member of the scientific advisory boards of Proxygen and Lyterian. The other authors declare no competing interests.

